# Structural and biochemical characterization of the mitomycin C repair exonuclease MrfB

**DOI:** 10.1101/2024.02.15.580553

**Authors:** Kelly A. Manthei, Lia M. Munson, Jayakrishnan Nandakumar, Lyle A. Simmons

## Abstract

Mitomycin C (MMC) repair factor A (*mrfA*) and factor B (*mrfB*), encode a conserved helicase and exonuclease that repair DNA damage in the soil-dwelling bacterium *Bacillus subtilis*. Here we have focused on the characterization of MrfB, a DEDDh exonuclease in the DnaQ superfamily. We solved the structure of the exonuclease core of MrfB to a resolution of 2.1 Å, in what appears to be an inactive state. In this conformation, a predicted α-helix containing the catalytic DEDDh residue Asp172 adopts a random coil, which moves Asp172 away from the active site and results in the occupancy of only one of the two catalytic Mg^2+^ ions. We propose that MrfB resides in this inactive state until it interacts with DNA to become activated. By comparing our structure to an AlphaFold prediction as well as other DnaQ- family structures, we located residues hypothesized to be important for exonuclease function. Using exonuclease assays we show that MrfB is a Mg^2+^-dependent 3’-5’ DNA exonuclease. We show that Leu113 aids in coordinating the 3’ end of the DNA substrate, and that a basic loop is important for substrate binding. This work provides insight into the function of a recently discovered bacterial exonuclease important for the repair of MMC-induced DNA adducts.

**GRAPHICAL ABSTRACT:** 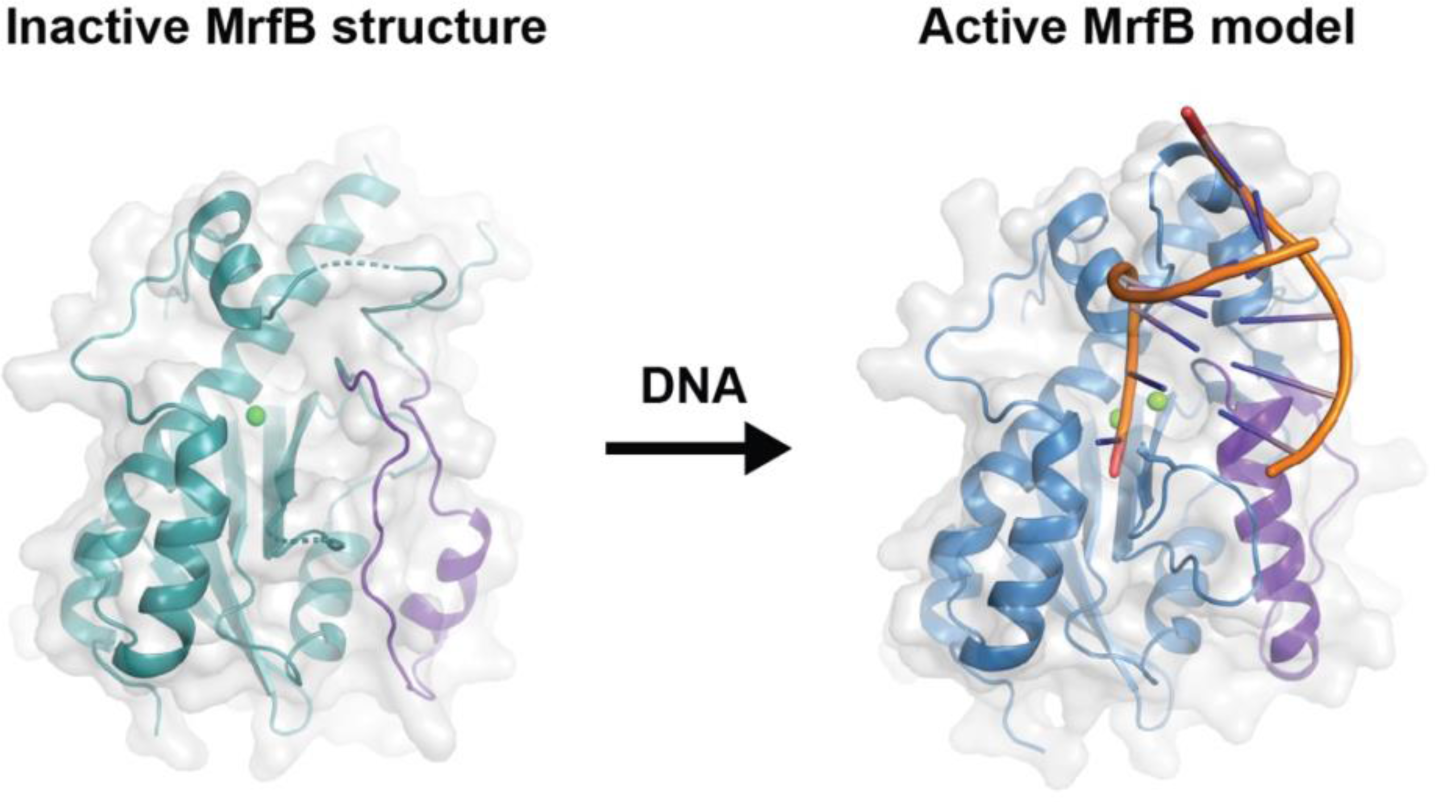

## INTRODUCTION

All organisms encode several exonucleases as part of their DNA repair and proofreading machinery to remove damaged or misincorporated nucleotides (1). Some organisms have evolved distinct machinery specific to the type of insults they frequently encounter. Soil-dwelling organisms have been known to produce a vast array of antimicrobial compounds, including the DNA damaging agent mitomycin C (MMC)(2). MMC was originally isolated from *Streptomyces caespitosus* and *S. lavendulae* and has subsequently been used as a chemotherapeutic (3). Once activated, it can react with guanine to form monoadducts, two adjacent guanines to form intrastrand crosslinks, or interstrand crosslinks if both strands have a CpG sequence (4–7). Intrastrand crosslinks and monoadducts are more common and can impede DNA replication or transcription, while interstrand crosslinks are deleterious because they prevent DNA from being unwound blocking replication and transcription (8–10). In bacteria the prevailing model is that MMC-induced DNA damage is repaired by the nucleotide excision repair (NER) pathway, however, interstrand crosslinks are proposed to require subsequent homologous recombination (8). NER is a general DNA repair pathway that responds to a wide range of bulky DNA adducts (11). In bacteria, UvrA initially scans the DNA alone or coupled to RNA polymerase to locate the DNA lesion, which then recruits the helicase UvrB to expose the lesion for UvrC to excise (8,12–14). Another helicase, UvrD, acts to release the damage and the lesion is filled in by DNA polymerase followed by ligation (15). For interstrand crosslinks, a double-strand break may result during repair which would require homologous recombination to complete repair (7).

We previously reported the discovery of Mitomycin C repair factor A (*mrfA*) and Mitomycin C repair factor B (*mrfB*), which encode a helicase and exonuclease, respectively (16,17). We showed that when *mrfA* and *mrfB* are deleted *B. subtilis* cells are specifically sensitive to MMC (16,17). We showed that MrfB is a metal-dependent DEDDh exonuclease, and that *mrfA* and *mrfB* are epistatic, suggesting that they work together to resolve the monoadduct and/or intrastrand crosslinks (17). The MrfAB pathway responds specifically to MMC insults and works independently of the UvrABC nucleotide excision repair pathway (17). While MMC-induced interstrand DNA crosslinks are lethal, the importance of MrfAB suggests that the more common monoadduct and/or intrastrand crosslink contribute to the overall MMC toxicity in *B. subtilis*. Therefore the MrfAB pathway allows *B. subtilis* to counteract MMC produced by competing soil bacteria (17). Recently the structure of MrfA was solved identifying a novel “rope skipping” mode of DNA unwinding providing important mechanistic insight into this newly discovered pathway (18). Although MrfA has been characterized biochemically, much less is known about the MrfB exonuclease.

MrfB is a member of the DnaQ superfamily of exonucleases with the DEDD motif, which includes the proofreading domains of many DNA polymerases, including *Escherichia coli* Pol I (*polA)* and the Pol III ε subunit (*dnaQ*) (17). The DnaQ superfamily also includes many standalone exonucleases, including *E. coli* ExoI and ExoX, mammalian TREX1/2, the exonuclease domain of the WRN helicase, as well as exoribonucleases such as RNase D, RNase T, NrnC, and Orn (19). The conserved structure for DnaQ/DEDD exonucleases contains a core mixed β-sheet containing 5 strands with conserved alpha helices surrounding the sheet. The first two DE residues are two amino acids apart on the first strand (β1) and the other two Asp are located on αB and αC (19). The DEDD residues coordinate two metals at the active site. There is a fifth conserved Tyr or His residue located four or five residues before the final Asp that is responsible for activating the nucleophilic water; in MrfB we showed this to be His258 (17). Thus, this family is further classified as DEDDh or DEDDy exonucleases, with MrfB residing in the DEDDh family (19). Aside from the catalytic exonuclease domain, at the C-terminus MrfB contains a tetratricopeptide repeat (TPR) domain, an α-helical repeating helix-turn-helix motif containing 34-amino acid repeats.

Based on sequence and structural models, the MrfB TPR fits the canonical domain containing three repeats capped with a C-terminal hydrophilic α-helix (AlphaFold entry P50837)(20–22). This domain packs into a superhelical shape where the inner concave surface can be used as a scaffold for protein or ligand interactions (20).

Here we sought to characterize MrfB both biochemically and structurally. We show that MrfB is a 3’-5’ DNA exonuclease and determined the apo structure of the exonuclease domain. Surprisingly, the structure appears to be in an inactive conformation, as a predicted α-helix was instead a random coil, which resulted in a key catalytic residue being displaced from the active site and unable to bind Mg^2+^. We then used structural comparisons to guide the generation of protein variants to evaluate exonuclease activity. Our results reveal the critical role of Leu113, conserved in many DEDD exonucleases as a wedge and in coordinating the 3’ end of the DNA substrate. We also examined several basic residues likely involved in DNA binding and showed their importance for exonuclease activity. Importantly, the basic residues and wedge were shown to be critical for *B. subtilis* to mitigate MMC damage in vivo. Together, our results identify several important features of the MrfB exonuclease as it protects *B. subtilis* from MMC-induced DNA damage.

## MATERIAL AND METHODS

### Protein Expression and Purification

All strains used in this study are listed in Table S2. *Full-length MrfB*. MrfB was expressed using pPB97 (10×His-Smt3-MrfB) and purified as previously described (16). MrfB variants were created by site-directed mutagenesis of the pPB97 plasmid, while truncations were created with Gibson assembly as detailed in the supplemental methods (Table S1)(23,24). For protein production, the appropriate plasmid was used to transform competent *E. coli* BL21(DE3) and plated on LB agar with 25 μg/mL kanamycin. Cultures were grown at 37°C with shaking at 200 rpm until the OD_600_ reached ∼0.6, the temperature was shifted to 20°C for 1 h, and expression induced with 0.5 mM IPTG overnight. Cell pellets were frozen at -80°C until purification, when they were resuspended in lysis buffer (50 mM Tris pH 7.5, 300 mM NaCl, 5% v/v glycerol), supplemented with 1 mM phenylmethanesulfonyl fluoride (PMSF) and protease inhibitor tablets (Pierce). Cells were lysed via sonication and clarified via centrifugation at 38,000 ×g for 30 min at 4°C (Sorvall SS-34 rotor). Clarified lysates were incubated with Ni^2+^-NTA-agarose pre-equilibrated in lysis buffer for ∼60 min, then spun (2000 rpm, 10 min), washed with lysis buffer, and poured into a gravity column. The column was washed extensively with lysis buffer, followed by 2 column volumes of wash buffer (50 mM Tris pH 7.5, 500 mM NaCl, 10 % v/v glycerol, 40 mM imidazole), and eluted with elution buffer (50 mM Tris pH 7.5, 150 mM NaCl, 10% v/v glycerol, 300 mM imidazole). The 10×His- Smt3 tag was removed by incubation with 20 μg mL^−1^ 6×His-Ulp1 and the addition of DTT to 1 mM at room temperature (RT) for 120 min. MrfB was dialyzed against 2 L of gel filtration buffer (25 mM Tris pH 7.5, 250 mM NaCl, 5% v/v glycerol) overnight at 4°C with stirring. The next day, the gravity column was re-used to bind 6×His-Ulp1, and MrfB eluted in the wash buffer with 40 mM imidazole. This wash was immediately loaded onto HiLoad Superdex 200-PG 16/60 column pre-equilibrated with gel filtration buffer (0.5 mL min^−1^). Peak fractions were pooled and concentrated using a 10 kDa Amicon centrifugal filter. Aliquots of 50-100 μL were flash frozen in liquid nitrogen at a concentration of ∼2-5 μM, and stored at -80°C. All proteins were purified twice, except for F171A which was unstable (Figure S1).

*Catalytic Core.* The MrfB catalytic core (residues 33-279) for crystallographic studies was purified with the following changes. Glycerol was removed from all buffers and 1 mM βME was added to the lysis and wash buffers. NaCl was increased to 250 mM to aid in solubility for the Ulp1 incubation. The dialysis and gel filtration buffer contained 25 mM Tris pH 7.5, 250 mM NaCl, and 1 mM TCEP. On the second Ni-NTA column, the catalytic core eluted in the flow through, was concentrated, and loaded to the HiLoad Superdex 200-PG 16/60 column pre-equilibrated with gel filtration buffer (0.5 mL min^−1^). Protein was used immediately for crystallization screens and the remainder flash frozen in aliquots.

### Crystallization and Structure Determination

#### Crystallization

Sparse matrix screens of the MrfB catalytic core at 10 g/L protein in 25 mM Tris pH 7.5, 125 mM NaCl, 1 mM TCEP, 5 mM MgCl_2_, 2 mM dTMP were set with a Crystal Gryphon (Art Robbins Instruments). Initial crystals were obtained via sitting drop vapor diffusion from the PEGs Suite (Qiagen) in 96 well plates and optimized with hanging drop vapor diffusion in 24 well plates. The condition that resulted in 2.1 Å diffraction was 1 μL of protein and 1 μL of well solution (0.1 M HEPES pH 7.5, 15% PEG Smear medium, 8% ethylene glycol). Crystals formed after two days and grew up to two weeks at 16°C. Crystals were harvested in 20 mM Tris pH 7.5, 125 mM NaCl, 1 mM TCEP, 5 mM MgCl2, 2 mM dTMP, 0.1 M HEPES pH 7.5, 20% PEG smear medium, cryoprotected with the addition of 25% ethylene glycol, and flash-frozen in liquid nitrogen.

#### Structure determination

X-ray diffraction data were collected at the Advanced Photon Source (LS-CAT beam line 21ID-D, λ = 1.127129) and were indexed and scaled using the DIALS User Interface within the CCP4i2 suite (25). The structure of the MrfB catalytic core was determined by molecular replacement using the AlphaFold model for the same residue range (22) as a search model in the program Phaser (26). The structure was manually inspected and fit in Coot (27) and refined using REFMAC5 Within CCP4i2 (28). Coordinate and structure factor files have been deposited in the Protein Data Bank (PDB ID code 8UN9).

### Exonuclease Assays

#### Urea-PAGE gel assay

20 nucleotide (nt) substrates were ordered from IDT and their sequences are listed in Table S1. The 20 nt DNA substrate oKM228 was ordered with an infrared (IR) dye on the 5’ end to visualize exonuclease activity. To test directionality, oJR348 contained an IR dye on the 3’ end, and to test exonuclease activity on RNA, oKM229 contained 11 nt of RNA at the 3’ end. 1 μM substrate in 20 mM Tris pH 8 was heated to 98°C for 1 min and allowed to cool RT prior to each reaction. Purified MrfB proteins were diluted to 1.67 μM in gel filtration buffer before being added to reactions to keep the NaCl concentration consistent. Final reaction conditions were 100 nM substrate, 0.5 μM MrfB, 75 mM NaCl, 20 mM Tris pH 8, 1 mM MgCl_2_ (or other metal as noted). After the appropriate time at RT, reactions were stopped with an equivalent volume of stop buffer (95% formamide, 20 nM EDTA, bromophenol blue). Stopped reactions were incubated at 98°C for 5 min, followed by cooling on ice. A ladder was generated using alkaline hydrolysis by incubating 500 nM of the RNA-containing oKM229 in 200 nM NaOH at 37°C for 10 min, then was stopped as above. Products were resolved using 20% denaturing urea-PAGE and visualized using the 800 nM channel of a LI-COR Odyssey CLx imager. Each reaction was repeated in triplicate, using at least two different protein preps for each variant or wild-type MrfB. The data were quantified using a band quantification macro for Fiji (dx.doi.org/10.17504/protocols.io.7vghn3w) (29). The intensity was divided by the intensity for the buffer control and converted to a percentage, then all replicates were plotted in GraphPad Prism version 10.0.3. Data are mean ± s.e.m. of three independent experiments. * 0.01 <P < 0.05, ** 0.001 <P < 0.01, *** 0.0001 < P < 0.001, **** p<0.0001 by one-way analysis of variance followed by Dunnett’s multiple comparisons post-test.

#### PicoGreen exonuclease assay

The PicoGreen (PG) assay was adapted from a published Bio-protocol and uses PG to monitor exonuclease activity on dsDNA (30). The PG dye in DMSO was stored at -20°C in 5 μL aliquots. Substrate stocks (Table S1, Figure S2) were annealed to a final concentration of 25 μM in PG annealing buffer (10 mM Tris, pH 8.0, 50 mM NaCl, 1 mM EDTA pH 8.0) by heating at 98°C for 5 min, then removing and wrapping the heat block in foil and allowing the reaction to come to RT, and finally freezing at -20°C. Streptavidin from Pierce (PI21122) was stored at 1 mg/mL at 4°C in nuclease-free water. For each reaction condition, the appropriate amount of substrate, 10X reaction buffer (200 mM Tris pH 8, 10 mM MgCl_2_) and streptavidin were first incubated for 15 min at RT. Then PG was added and the reaction was aliquoted into a black 96-well plate (Greiner 655209) and incubated at 30°C in the plate reader for 15 min. MrfB variants were diluted to 1.67 μM in gel filtration buffer at RT, then added to the appropriate well (or gel filtration buffer alone) in duplicate for each reaction. Timepoints were taken every 90 s for 2 h at 30°C on a Tecan Infinite M1000 plate reader with a 3 s shake before each read (483 nm excitation, 530 nm emission). The data were processed in GraphPad Prism version 10.0.3 as described in the Bio-protocol, using known dsDNA substrate lengths of 80, 60, 40, and 20 bp to create a calibration curve (30). A negative control of a dsDNA 80 bp substrate with blocks on all 4 ends and wild-type MrfB was used to correct for photobleaching and background by dividing the data by the average reads from the negative control (Figure S2). Each reaction was an average of two technical repeats done in triplicate, using at least two protein preps for each variant. Each reaction was fit in GraphPad Prism with a one-phase decay and the rates were analyzed statistically as compared to WT with one-way analysis of variance followed by Dunnett’s or Tukey’s multiple comparisons test in GraphPad Prism.

### Spot Titer Assays

#### Plasmid and strain construction

Plasmids were created with either site-directed mutagenesis or Gibson assembly as detailed previously and in the supplemental methods (16,17,24). *Pxyl- mrfB* variant plasmids were used to transform *B. subtilis* Δ*mrfB* strain using double crossover recombination at the *amyE* locus. Replacement of the *amyE* locus was verified by the inability to use starch and by diagnostic colony PCR.

#### Spot titer sensitivity assays

*B. subtilis* strains were struck out onto the appropriate LB agar plates and incubated at 37°C overnight. A single colony per strain was inoculated in 2 mL LB media in 14 mL round bottom culture tubes, supplemented with 5 µg/mL chloramphenicol as needed. Cultures were incubated on a rolling rack at 37°C until the OD_600_ was 0.5-0.8.

Cultures were normalized in 200 µL to an OD_600_ of 0.5 using 0.85% w/v sterile saline and were serially diluted in saline to 10^-5^. Then 5 µL of the serial dilutions were spotted onto LB agar plates and grown at 30°C for 15 hours. All spot titers were performed with two duplicates in biological triplicate and were imaged using the white light source in the Alpha Innotech MultiImage Light Cabinet, with exposure, brightness, and contrast edited in Adobe Photoshop.

## RESULTS

### MrfB is a 3’-5’ DNA exonuclease

We previously showed that MrfB is an exonuclease that can degrade both linear and nicked dsDNA in the presence of Mg^2+^ (17). To understand MrfB metal dependence we tested various divalent metal ions assayed at 1 mM and observed that MrfB was active in the presence of Mg^2+^ or Mn^2+^ (Figure 1a). As free Mg^2+^ and Mn^2+^ are at concentrations of ∼1 mM and ∼10 µM in vivo, respectively, we used 1 mM Mg^2+^ in the subsequent experiments (31–33). We tested the directionality of the nuclease activity using 20 nt substrates with either a 3’ or 5’ end label. Exonuclease activity was only observed when the label was on the 5’ end, indicating that MrfB degrades DNA in the 3’–5’ direction (Figure 1b). A third substrate containing 11 nt of RNA on the 3’ end was used to determine if MrfB was specific for DNA. Very little activity was observed for the 3’ end RNA substrate. With these results we conclude that MrfB is a 3’–5’ DNA exonuclease (Figure 1a,b).

**Figure 1.**
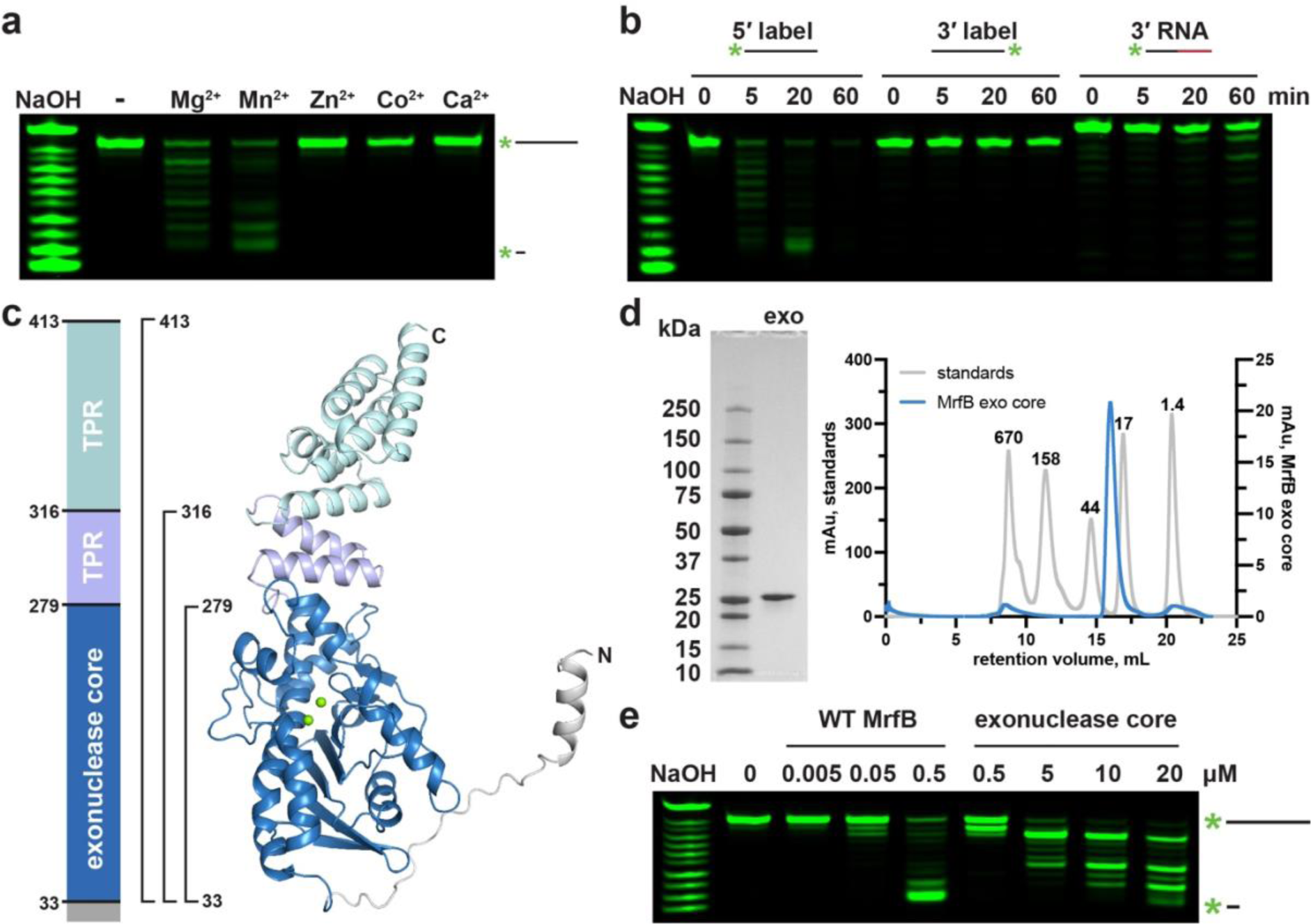
Exonuclease assays with 5’ labeled DNA confirm directionality and metal-dependence. (a-b) Initial biochemical assays performed with full-length WT MrfB. All assays used 20-mer oligos with various labels as depicted above. NaOH refers to the 3’-labeled RNA substrate treated with alkaline hydrolysis for a ladder. (a) Assays stopped at 20 min with 1 mM of the indicated metal show that MrfB can use Mg^2+^ or Mn^2+^. (b) Comparison of 20-mer oligos labeled on different ends and a third with RNA at the 3’ end showing that, MrfB is a 3’-5’ exonuclease with a preference for DNA. (c) Left: domain architecture of full length MrfB. TPR stands for tetratricopeptide repeat. Right: AlphaFold model of full-length MrfB (entry P50837), showing the three truncation constructs that were cloned and purified with brackets. (d) Purified exonuclease core (residues 33-279, 28.6 kDa) electrophoresed on an SDS-PAGE gel (left), and on a Superdex 200 Increase 10/300 GL column on right (blue trace). The gray traces show standards with their molecular weight indicated above each peak, confirming that the exonuclease core is a monomer. (e) The same assay as in (a) with increasing concentrations of full-length MrfB and the exonuclease core.

### The structure of the MrfB catalytic core appears inactive

We sought to determine the structure of MrfB using X-ray crystallography and gain insight into its mechanism. We initially purified full-length MrfB from overexpression in *E. coli* but its yield was poor and the protein was unstable at high concentrations. We then used an AlphaFold model (entry P50837 (22)) to design truncations of MrfB to increase stability following overexpression (Figure 1c). As the N-terminal domain was predicted to be unstructured, we proceeded with constructs lacking the first 32 amino acids. Three constructs were examined: residues 33-413, which removed only the N-terminus, 33-316, which removed all but the last two α-helices comprising the first TPR repeat, and 33-279, which comprised the core exonuclease domain (Figure 1c). The exonuclease core domain, which displayed the highest expression level, was subjected to crystallization trials (Figure 1d, S3). The catalytic core was confirmed to have exonuclease activity, although the activity was approximately tenfold reduced compared to the WT enzyme on a 20 nt ssDNA substrate (Figure 1e). The decreased activity observed with the catalytic core agrees with our previous in vivo findings showing the *ΔTPR* construct from residues 1-284 was able to complement *ΔmrfB*, but only at the highest induced expression tested (17).

The MrfB exonuclease core was purified and initial sparse matrix screens were set with the addition of MgCl_2_ and dTMP, which would represent the nucleotide product of the exonuclease reaction (34). Multiple hits were obtained and refined, with one crystal diffracting to 2.1 Å. The structure was solved by molecular replacement with the AlphaFold model for residues 33-279 as a search model (Table 1, Figure 2a-b). Two molecules of MrfB were present in the asymmetric unit. The dimeric status in the crystal likely arises from crystal packing as the exonuclease core eluted as a monomer on gel filtration (Figure 1d). Density was observed for residues 34-111, 119-202, and 213-279. There was no density observed for dTMP, and only one Mg^2+^ (site A) was observed at the active site, coordinated by residues Asp107, Glu109, and Asp262. Furthermore, a predicted α-helix from residues 171-183 was in a random coil, which resulted in Asp172 being displaced away from the active site and unable to coordinate a Mg^2+^ ion in the B site (Figure 2a-b). Because the mechanism for DnaQ exonucleases coordinates two metal ions, we expect that this structure represents an inactive state, and the AlphaFold model likely represents an active state (35,36). We propose that MrfB resides in an inactive state in the absence of DNA. Upon interaction with DNA, the region spanning Asp172 folds into a helix to recruit the site B Mg^2+^ ion, establishing the active site geometry and activating the exonuclease (Movie 1).

**Figure 2.**
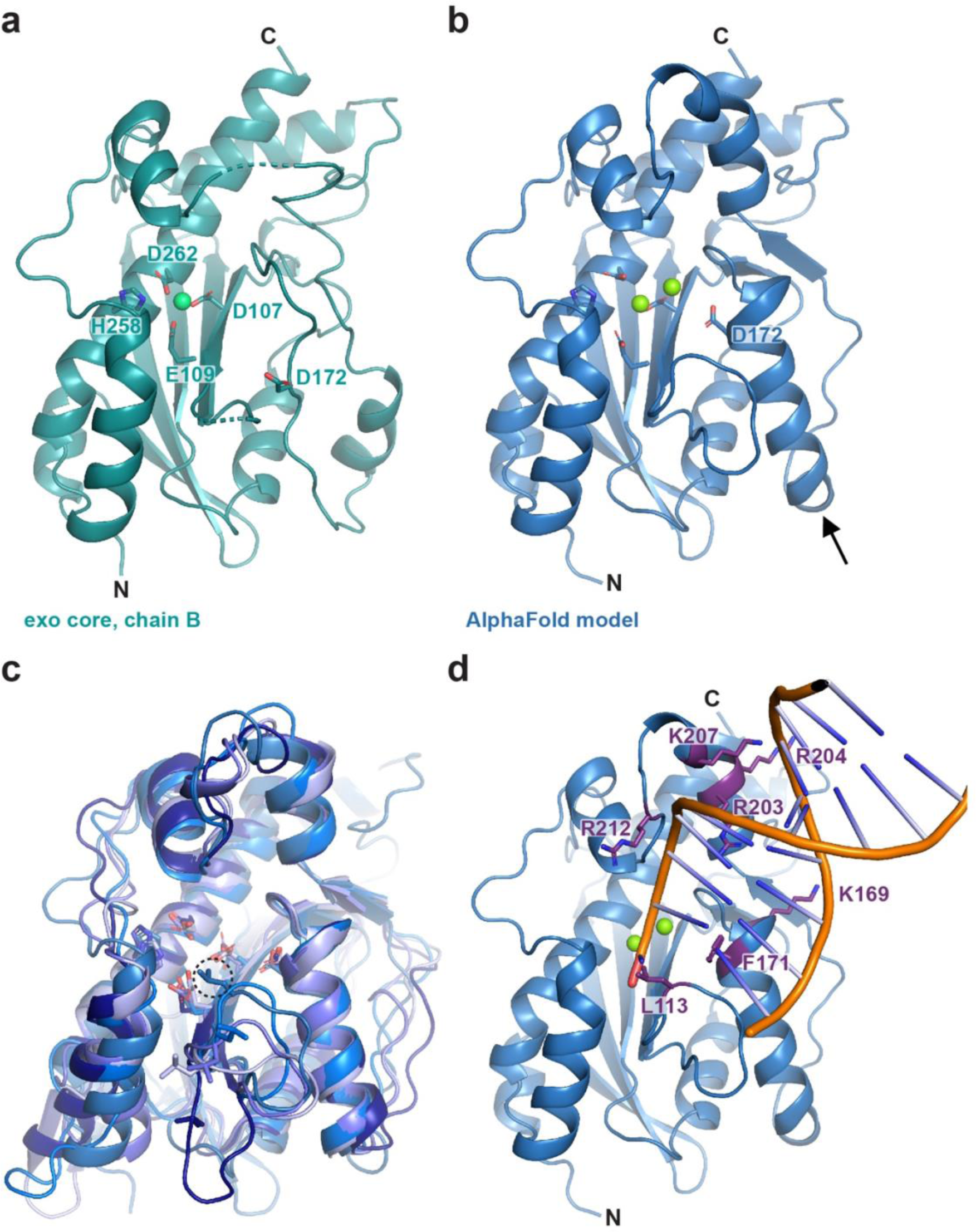
**The structure of the MrfB exonuclease core**. (a) The 2.1 Å structure appears to be an inactive state due to an unwound helix that pulls D172 away from the active site, and likely leading to only one observed Mg^2+^ ion (green sphere). DEDDh residue side chains are shown as sticks. Density was not observed for loops spanning residues 112-118 and 204-212 (dashed lines). (b) An AlphaFold model of residues 33-279, modeled using the Colab notebook that uses no homologous templates (https://colab.research.google.com/github/deepmind/alphafold/blob/main/notebooks/Alpha Fold.ipynb) (22). The AlphaFold model is more consistent with an active state, with a helix containing Asp172 spanning residues 171-183 (see arrow), which positions all DEDDh residues at the active site. Mg^2+^ ions are modeled from PDB 5YWU. (c) A consistent model for the exonuclease core is observed from different software, aside from the loop containing Leu113. Models are from AlphaFold (Leu113 in dashed circle), I-TASSER and SWISS-MODEL (22,37,38). (d) The AlphaFold model is consistent with other exonuclease structures that position Leu113 at the active site. The DNA is modeled from alignment with an ExoX structure (PDB 4FZZ)(40). Potential DNA-interacting residues are shown with side chains as sticks.

**Table 1.** Data collection and refinement statistics.

|  | **MrfB exo core** |
| --- | --- |
| **Data collection** |  |
| Space group | P1 2<sub>1</sub> 1 |
| Cell dimensions |  |
| *a*, *b*, *c* (Å) | 72.87, 37.93, 92.62 |
| $\alpha$, $\beta$, $\gamma$ (°) | 90.00, 99.15, 90.00 |
| Resolution (Å) | 45.72 – 2.10 (2.16 – 2.10) |
| *R*<sub>merge</sub> | 0.064 (0.489) |
| *I* / σ*I* | 14.2 (2.7) |
| Completeness (%) | 99.6 (99.3) |
| Redundancy | 6.5 (6.0) |
| CC<sub>1/2</sub> | (0.917) |
| **Refinement** |  |
| Resolution (Å) | 45.72 – 2.10 |
| No. reflections | 29515 (2294) |
| *R*<sub>work</sub> / *R*<sub>free</sub> | 0.175/0.232 |
| No. atoms |  |
| Protein | 3766 |
| Ligand/ion | 2 |
| Water | 241 |
| *B*-factors (Å<sup>2</sup>) |  |
| Protein | 55.27 |
| Ligand/ion | 47.89 |
| Water | 61.05 |
| R.m.s. deviations |  |
| Bond lengths (Å) | 0.0084 |
| Bond angles (°) | 1.41 |
| Ramachandran statistics (%) |  |
| Favored | 98.21 |
| Allowed | 1.79 |
| Outliers | 0.0 |
| MolProbity score | 1.02 |
| Clashscore, all atoms | 2.41 |

To further characterize the determinants of MrfB exonuclease activity, we designed a set of protein variants. First, the catalytic DEDDh residues were each mutated to alanine and purified, creating D107A, E109A, D172A, H258A, and D262A. To identify other potentially important residues, the AlphaFold model was also compared to other models generated in I-TASSER (37) and SWISS-MODEL (38) (Figure 2c). While these models agree on the overall fold and orientation of the DEDDh residues, the loop from residues 111-120 has high variation between models. This information combined with the lack of density observed in our structure suggests that this loop is dynamic. Leucine 113 is conserved in other exonucleases and has been shown to act as a wedge to break the terminal base pairing and coordinate the 3’ end of the DNA (34,39–41). In the AlphaFold model Leu113 is positioned to act as a wedge (Figure 2c). Thus, we engineered the L113A mutation to investigate its effect on MrfB activity (see below).

To identify residues important for DNA binding, we compared our structure and the AlphaFold model to DNA-bound structures of similar exonucleases, such as ExoX (40). From this analysis, we substituted basic residues poised to interact with DNA, creating K169A, R203A, R204A, K207A, R212A, and a basic loop mutation that contained R203A, R204A, K207A, and R212A (RRKR) (Figure 2d). In the AlphaFold model, Phe171 is poised to base stack with the DNA, and in our structure is instead flipped into a hydrophobic pocket. We hypothesized that the movement between the two structures could be a mechanism to protect this hydrophobic residue from solvent in the absence of DNA. However, F171A protein yields were very low, suggesting that this mutant is unstable and was not pursued further.

### Several MrfB variants show reduced exonuclease activity on ssDNA

We purified the MrfB variants described above and assayed for exonuclease activity on a ssDNA substrate (Figure S1). The MrfB variants were initially examined in a gel-based assay with a 20 nt ssDNA substrate. As expected, the DEDDh catalytic point variants had decreased activity (Figure 3a). D107A, E109A, and H258A had significantly lower activity, while D262A retained some activity and was not statistically different from WT MrfB (Figure 3a). Asp107, Glu109, and Asp262 coordinate the Mg^2+^ (site A) in the structure, while His258 acts to activate the nucleophilic water. D172A also retained some activity, which agrees with our previous report that D172A could complement Δ*mrfB* in vivo at the highest induced expression tested (17). Asp172 is out of position in our structure as described above but is expected to coordinate the site B Mg^2+^ along with Asp107.

**Figure 3.**
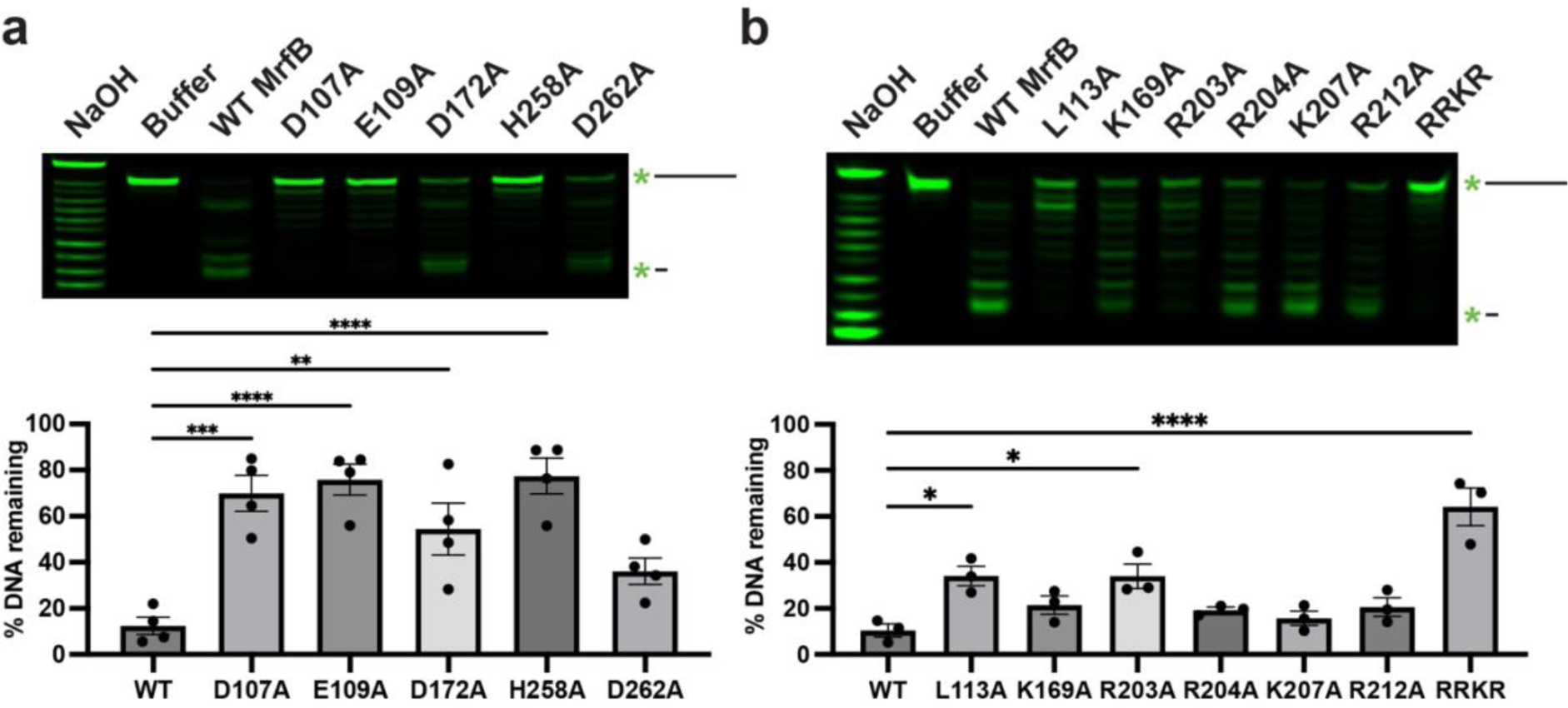
**MrfB variant exonuclease activity on ssDNA**. (a) DEDDh variants reveal that D107, E109, and H258A are required for catalytic activity. (b) DNA-interacting variants show the importance of the basic loop and Leu113. (a-b) Both panels show a gel-based assay with 5’ labeled DNA on top, and quantification of the remaining DNA from the top of the gel at the bottom as compared to the buffer control (see Methods). In all assays exonuclease activity is measured using a 20-mer ssDNA substrate. Data are mean ± s.e.m. of 3-4 independent experiments. * 0.01 <P < 0.05, ** 0.001 <P < 0.01, *** 0.0001 < P < 0.001, **** p<0.0001 by one-way analysis of variance followed by Dunnett’s multiple comparisons post- test.

Many of the other single point variants designed to disrupt DNA binding retained activity that was not statistically different from WT, including K169A, R204A, K207A, and R212A (Figure 3b). However, R203A had a significant loss in activity (10% remaining substrate DNA for WT vs. 34% for R203A) and the basic loop RRKR had a further reduction in activity with 64% of the DNA remaining. These results indicate that while one single point mutation to the basic loop is mostly not deleterious, mutating multiple residues has a detrimental effect on exonuclease function. Finally, the L113A variant also had a significant decrease in activity (34% remaining DNA), which indicates that Leu113 likely acts in part to orient the 3’ end of the DNA in the active site.

### L113A and RRKR MrfB variants show defects in exonucleolytic processing of dsDNA

We used a PicoGreen fluorescence assay to quantitively assess exonuclease activity on several dsDNA substrates (30) (Figure S2). MrfB variants were first analyzed with a blunt 80 bp dsDNA substrate that was blocked on one side with biotin-streptavidin, so that only one free 3’ end was available (Figure S2b). For the DEDDh variants, D107A, D172A, and D262A were analyzed, and all showed no exonuclease activity over the two-hour incubation period with DNA (Figure 4a). As in the gel-based assay, most single point mutants to potential DNA-binding residues did not affect activity, and in this assay R203A also had WT activity. L113A had a significantly decreased rate that appeared to be biphasic with a slower initial phase. This result is consistent with a critical role for Leu113 in orienting DNA in the active site and possibly acting as a wedge to help separate dsDNA as it moves into the active site. Finally, the RRKR basic loop mutation showed no activity, similar to the gel-based assay, emphasizing its important contribution to DNA-binding.

**Figure 4.**
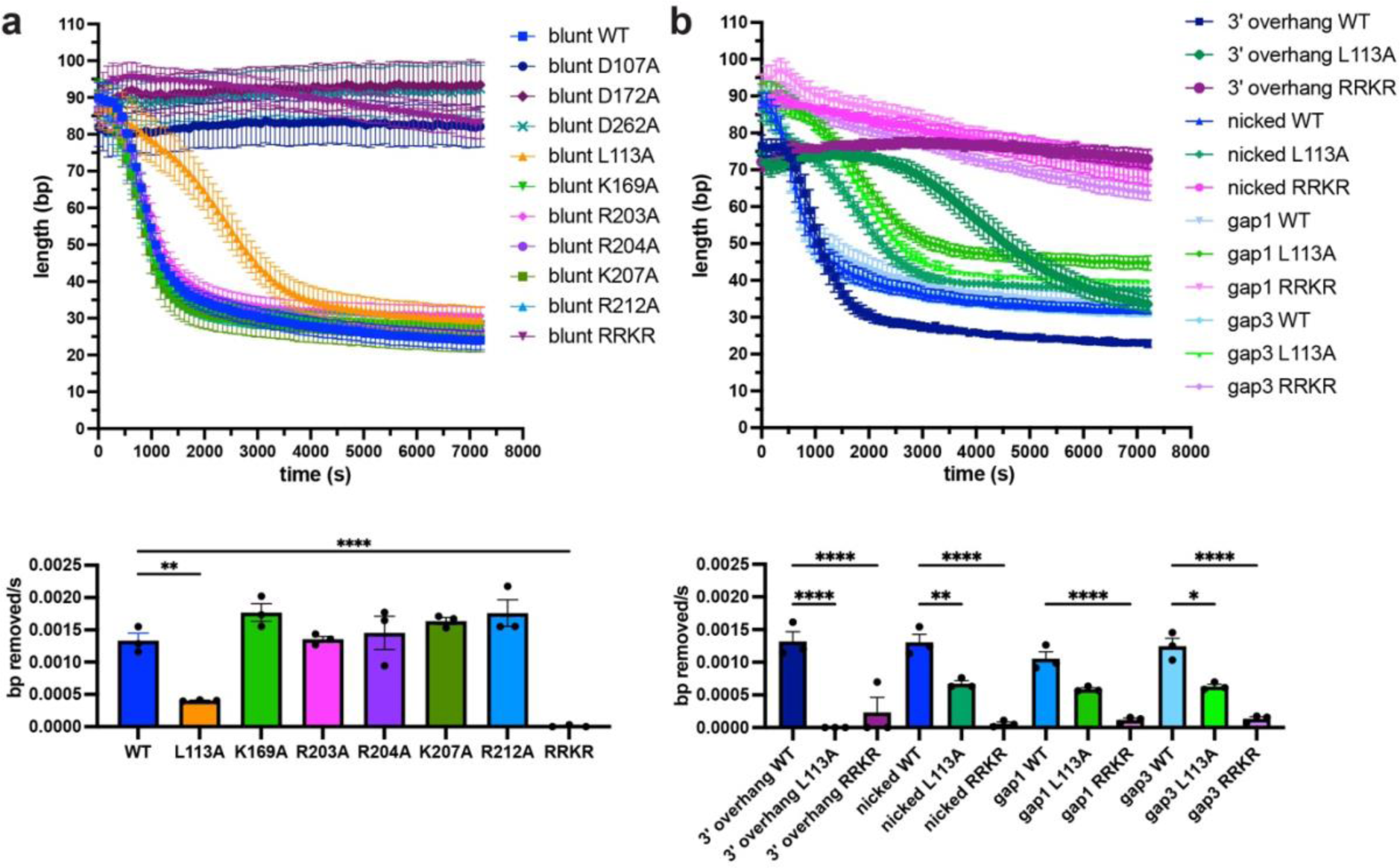
PicoGreen assay examining MrfB variants on various dsDNA substrates. Fluorescence was monitored for 2 h to examine nuclease activity on (a) an 80 bp blunt or (b) a 3’ overhang, nicked, and gap of 1 or 3 nucleotide substrate (see Figure S2 for substrates). The data in the top panels were fit to a one phase decay model to determine the rate, which was then compared in the bottom panels. Data are mean ± s.e.m. of 3 independent experiments. * 0.01 <P < 0.05, ** 0.001 <P < 0.01, *** 0.0001 < P < 0.001, **** p<0.0001 by one-way analysis of variance followed by (a) Dunnett’s or (b) Tukey’s multiple comparisons post-test.

**Figure 5.**
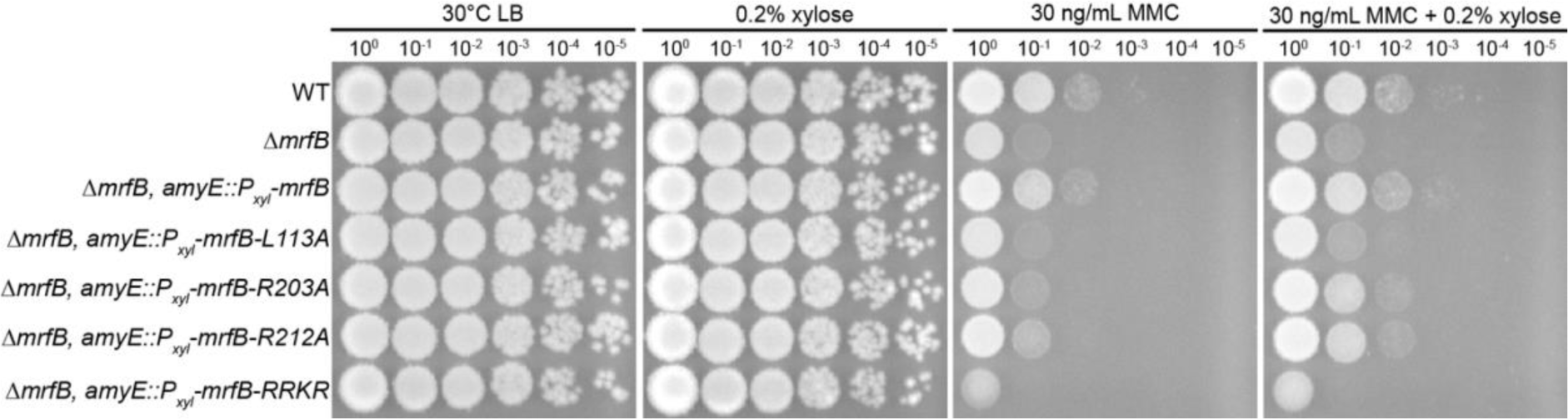
The *mrfB L113A* and *RRKR* are unable to complement Δ*mrfB* in vivo. The addition of MMC is more deleterious in a Δ*mrfB* background, with growth restored by complementation with WT, and partially restored by complementation with *R203A*, and *R212A mrfB* when overexpressed with xylose. The *L113A* and *RRKR* mutants are unable to complement *ΔmrfB* even when induced with xylose.

We analyzed WT, L113A, and RRKR with a panel of substrates including a nicked substrate, gaps of 1 and 3 nt, and a 3’ overhang of 15 nt to determine if MrfB shows a DNA substrate preference (Figure 4b, S2). WT MrfB and the variants degraded the DNA substrates with similar activity, except for L113A, which had virtually no activity on the 3’ overhang.

Further, L113A showed a longer initial lag phase with the 3’ overhang substrate than other substrates. This could indicate that L113A has more difficulty degrading a substrate with a ssDNA or a ssDNA/dsDNA junction as the substrate has a 15 nt overhang. Based on these data, we hypothesize that Leu113 not only acts as a wedge, as observed for *E. coli* ExoX, but is also similar to the eukaryotic TREX1 where the leucine was shown to be important for activity on both ssDNA and dsDNA (40,42). If Leu113 acted solely as a wedge in MrfB, we would have expected preferred degradation of ssDNA. Our data suggest that Leu113 is also critical to coordinate the 3’ end at the active site.

### Leu113 and the basic loop are important for DNA repair in vivo

To examine the importance of Leu113 and several other residues shown to be important for activity in vitro, we assayed for a phenotype corresponding to repair of MMC-induced DNA damage in vivo. We ectopically expressed *L113A, R203A, R212A*, and the *RRKR* basic loop variants in a *mrfB* deletion background as described (17). We previously showed that the catalytic DEDDh point mutants could not complement the *ΔmrfB* sensitivity to MMC. Here we show that *L113A* and *RRKR* also failed to complement the *ΔmrfB* phenotype, while *R203A* and *R212A* were able to restore growth to near WT levels. These results parallel the biochemical assays shown above and further support the MrfB activity measured *in vitro*. We conclude that Leu113 and the RRKR basic loop are critical for MrfB function in removing MMC damage in vivo.

## DISCUSSION

Mitomycin C induces DNA damage by reacting with guanines to form monoadducts, intrastrand and interstrand crosslinks (7,43). We have previously shown that the MrfAB pathway works in addition to canonical NER to remove the monoadduct and/or intrastrand crosslinks (17). Here we have characterized the MrfB exonuclease structurally and biochemically, revealing biochemical results that are supported by mutant phenotypes in vivo. Unexpectedly, the structure of the MrfB exonuclease core appears inactive, due to the lack of a predicted α-helix from residues 171-183, which resulted in Asp172 being displaced away from the active site and unable to coordinate a site B Mg^2+^ (Figure 1). It is possible that this conformation was induced by crystal packing, and indeed there are symmetry mates nearby, but it is difficult to speculate that this alone would induce such a striking change in conformation (Figure S4). The inactive conformation could also be due to the smaller exonuclease core that was crystallized, however, in the AlphaFold model of full-length MrfB the rest of the protein does not appear to contact the exonuclease domain especially near the α-helix. Instead, we propose this conformation protects hydrophobic residues of MrfB from solvent when it is not engaged with DNA and/or MrfA, and becomes activated by those interactions. Lending further support to the dynamic nature of this region, the highest B-factors of the structure are observed between residues 174-189 (Figure S4c). Additionally, multiple AlphaFold models displayed varying degrees of confidence in that region, with some even pointing Phe171 inward (Figure 6a-b).

**Figure 6.**
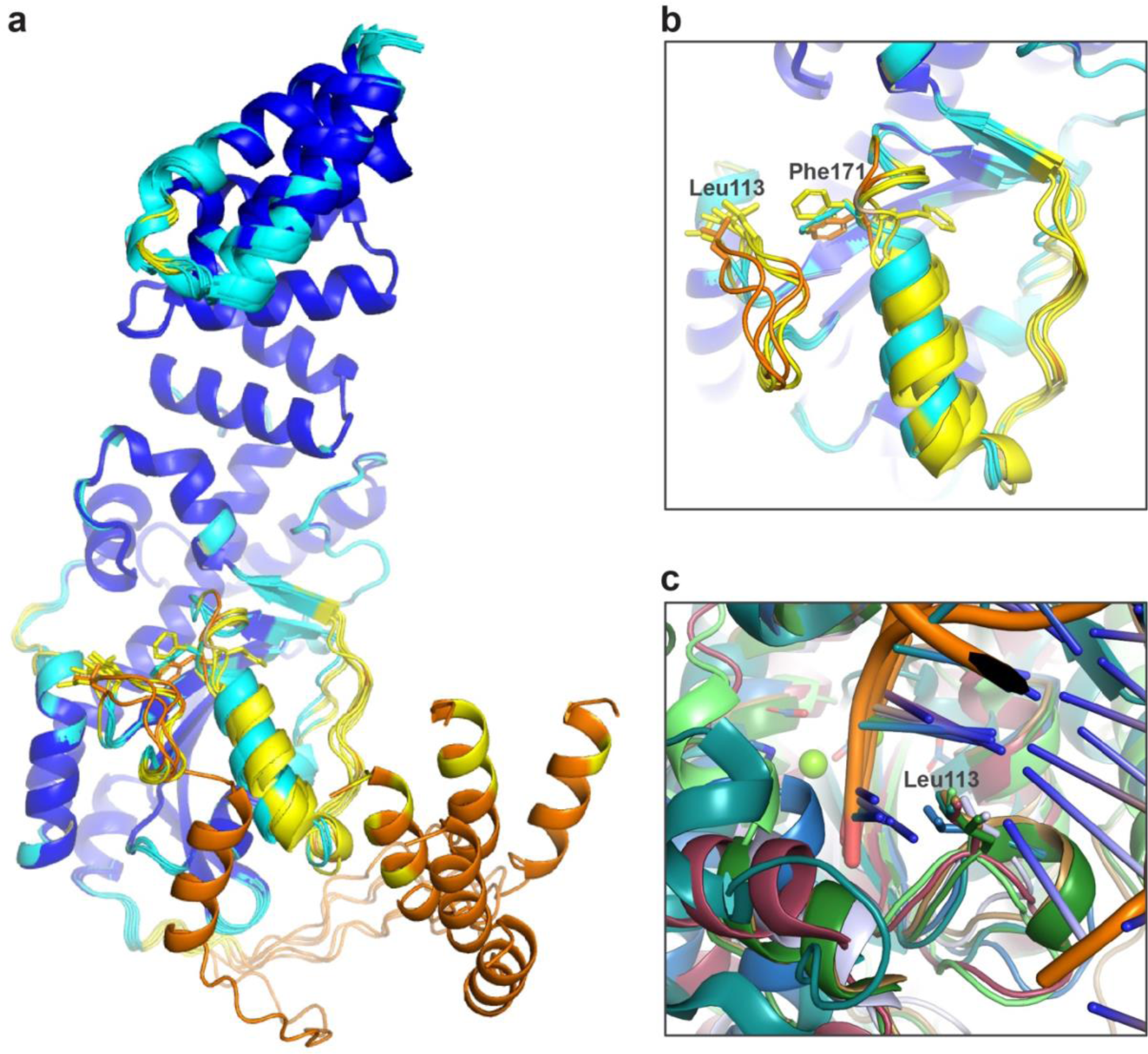
MrfB structural modeling and comparisons. (a-b) A comparison of the deposited AlphaFold model with 5 models determined without a structural template, showing some uncertainty for the α-helix preceding Phe171 and Asp172. (22). AlphaFold models are colored by pLDDT, a model confidence score. Very high confidence (>90) scores are dark blue, high confidence is light blue (70–90), low scores are yellow (50–70), and very low scores are orange (<50). (c) Different crystal structures aligned at the active site to show a variety of DEDD exonucleases use a leucine wedge at the active site. Structures shown: MrfB AlphaFold model, blue; ExoX (PDB 4FZZ) light blue; TREX2 (PDB 6A47) tan, TREX1 (PDB 2O4I) dark green; Klenow fragment (PDB 1KLN), teal; NrnC (PDB 7MPO), light green (22,34,39–41,50).

During the conformational change that results in MrfB activation, residues Asn167 and His195 are at the boundaries of the movement, with both residues flipping conformation (Movie 1). The region between those residues must rotate for the helix to form, and while the hydrophobic nature of the interior is maintained, Phe171 gets flipped out towards the active site. In the active state, Phe171 is able to interact with DNA and Asp172 is able to coordinate the site B Mg^2+^. Both residues are conserved in the Exo II motif (44). Our inability to purify the F171A variant also suggests that it has a critical role in protein stability, which appears to be important to the inactive state because it is pointed in a hydrophobic pocket. However, after Phe171 is flipped out, it is poised to base stack the penultimate deoxyribose of the

DNA substrate, which is consistent with the role of the conserved Tyr or Phe in many other DEDD exonucleases (36,45). Indeed, mutation of Tyr423 to Ala in the Klenow fragment severely disrupted exonuclease activity on ss and dsDNA, and Tyr423 was also proposed to contribute in duplex melting (45). However, there are many apo structures of DEDD exonucleases that have this conserved Phe/Tyr pointed towards the active site, and no other similar structural rearrangements have been observed (46–50).

While our structure does not contain DNA, we used comparisons to other DEDD exonucleases to identify residues critical for DNA binding. A common feature is the use of a leucine residue to act as a potential wedge and/or in coordination of the 3’ end (Figure 6c). In ExoX mutations to Leu12 and Gln13 were shown to be specifically important for dsDNA exonuclease activity, and were hypothesized to act as a wedge to break the duplex and feed the 3’ end into the active site (40). A similar activity profile was observed for the Klenow fragment mutant L361A, and Leu361 was expected to interact with the 3’ nucleotide (45,51).

The eukaryotic TREX1 and 2 utilize a small helix containing leucine, and mutations in TREX1 were shown to affect activity on both ssDNA and dsDNA, similar to MrfB (41,42). Finally, a leucine wedge is also used in some exoribonucleases such as Orn and NrnC (50,52). Here, our data strongly support the orientation for Leu113 observed in the AlphaFold model where it is poised to melt duplex DNA and interact with the 3’ end. In our biochemical analysis, L113A had decreased activity on all DNA substrates examined, and struggled to move through the ssDNA in the 3’ overhang substrate (Figure 3b, 4). The L113A variant did not fit a one phase decay model and instead appeared to have a slower initial phase (Figure 4). This points to the importance of this residue in not only melting the duplex, but also in coordinating the 3’ end since it affects both ssDNA and dsDNA degradation.

Another common feature of DEDD exonucleases are the basic loop residues to coordinate substrate binding. We used comparisons to ExoX to design point mutants that we expected would specifically affect substrate or complementary strand binding (40). However, the single alanine substitutions had similar activity to WT MrfB, aside from R203A on ssDNA, which indicates a role in coordinating the substrate strand (Figure 3b). When the basic loop alanine substitutions were combined to create the RRKR variant, no exonuclease activity remained, indicating that these residues likely interact with DNA in a coordinated effort where one change is not deleterious. One caveat of our study is that we have not yet been able to examine the expected substrate containing an MMC adduct. This perhaps explains the slow rate and high concentrations needed in biochemical assays, and it remains possible that the difference between mutants might become more apparent on an MMC-bearing substrate. MrfA might also be required to stabilize and promote an active conformation both in vitro and in vivo.

While we have utilized other DEDDh DNA exonuclease structures such as ExoX and TREX1 to aid our biochemical analyses, we also compared MrfB to other exonucleases with similar folds based on the top matches using the DALI web server (53). In all the structures examined, the same core fold is present consisting of β1-β2-β3-αA-β4-αB-β5-αC as well as a helical region that often contains substrate-binding residues between β5 and αC (Figure 7). The top structural similarities are the proofreading exonuclease domains from archaeal B-family polymerases (PDB 4FLU), yeast Pol ε (PDB 6WJV, 6S2E), an inactive exonuclease domain from human POLA1 (PDB 8D0B, 7OPL), and Gram-negative bacterial NrnC (PDB 7MPO) (50,54–58). Aside from the inactive POLA1, these are DEDDy exonucleases, with the Tyr being part of a longer αC, whereas the His in DEDDh is typically found in a loop preceding αC (Figure 7a-b). As similar grouping has been described for the exoribonucleases NrnC/RNaseD that are DEDDy compared to Orn/RNase-T which are DEDDh (50). MrfB has the longer αC and is structurally more similar to the DEDDy exonucleases, yet is DEDDh, and also contains a long N-terminal insertion prior to β1 (Figure 7c). Thus, MrfB appears to be structurally distinct from other known DEDD structures. Future work will be important to determine the active state with metals and DNA to improve our mechanistic understanding of this class of exonucleases.

**Figure 7.**
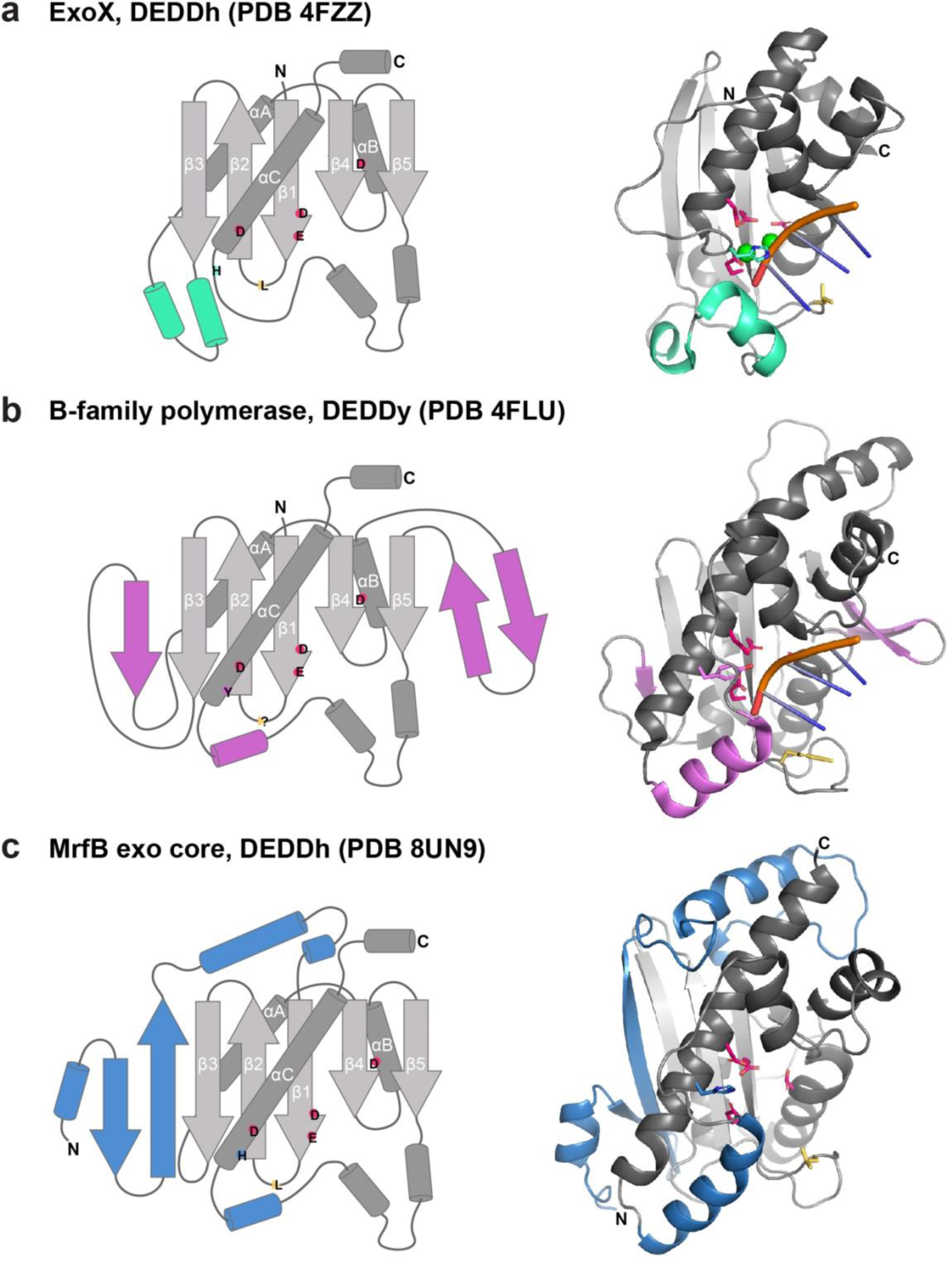
DEDD exonuclease structural comparisons. (a-c) In all 3, the fold topology is shown on the left, and structural features colored the same on the right. The canonical fold is in gray. DEDDh/y residues are shown with the single letter in their approximate location, and a yellow wedge for any leucine or other potential wedge location. In the structural models, the same highlighted active site residues are shown with sticks. (a) ExoX, PDB 4FZZ, DEDDh (40). 3 bases of the 3’ end are shown for clarity, but the structure contains dsDNA. Green spheres are Na^+^. (b) The proofreading exonuclease domain from *Pyrococcus abyssi* B-family polymerase bound to dsDNA, which is melted so only ssDNA is at the exonuclease active site (54). There is no need for a leucine wedge as the pink β-hairpin stabilizes the other strand of the duplex and only ssDNA is fed into the active site. (c) MrfB exo core as described in this paper, with part of the N-terminal insertion starting at residue 33 shown in blue.

Here we have focused on the exonuclease core, however the role of the TPR domain remains unknown. TPR domains are known to act as a scaffold, and while it is tempting to suggest this domain interacts with MrfA, our previous bacterial two-hybrid data suggest MrfA interacts with the exonuclease core (17,20,59). If these proteins indeed interact it will be important to determine the interface and if the interaction is ATP- and/or DNA-dependent.

There are many basic residues on the surface of the TPR, suggesting it could be important for DNA interactions. It could also interact with another unknown protein contributing to the repair of MMC-induced adducts. In summary, we have revealed the structure of an inactive exonuclease core of MrfB and described the residues important to MrfB function. Future studies will be directed toward understanding how MrfA, MrfB, and any other unknown partners coordinate their efforts during MMC repair.

## DATA AVAILABILITY

Coordinate and structure factor files have been deposited in the Protein Data Bank (PDB ID code 8UN9).

## AUTHOR CONTRIBUTIONS

This study was conceived and designed by KAM and LAS. Experiments were performed by KAM and LMM. KAM performed the experiments shown in Figures 1-4 and 6, LMM performed the experiments shown in Figure 5. Data analysis was performed by KAM, LMM, JN and LAS. The manuscript was written by KAM and LAS. The manuscript was revised by KAM, LMM, JN and LAS.

## ACKNOWLEDGEMENTS

We would like to thank Dr. Valerie Tesmer for assistance with X-ray crystallography, Dr. Randy Stockbridge for the use of the Crystal Gryphon, and Thomas Svetich for assistance in creating the *B. subtilis* strains. This research used resources of the Advanced Photon Source, a U.S. Department of Energy (DOE) Office of Science User Facility operated for the DOE Office of Science by Argonne National Laboratory under Contract No. DE-AC02-06CH11357. Use of the LS-CAT Sector 21 was supported by the Michigan Economic Development Corporation and the Michigan Technology Tri-Corridor (Grant 085P1000817).

---

**Movie 1. Movie depicting morph from inactive structure to active model.** Active site residues are shown in sticks, and Phe171 in pink sticks to highlight its conformational change. The greatest movement occurs from residue 167-195, shown in purple. The movie was rendered in PyMOL (The PyMOL Molecular Graphics System, Version 2.5.8 Schrödinger, LLC).

## REFERENCES

1. Friedberg, E.C., Walker, G.C., Siede, W., Wood, R.D., Schultz, R.A. and Ellenberger, T. (2006) DNA Repair and Mutagenesis: Second Edition. American Society for Microbiology, Washington, DC.

2. Procopio, R.E., Silva, I.R., Martins, M.K., Azevedo, J.L. and Araujo, J.M. (2012) Antibiotics produced by Streptomyces. Braz J Infect Dis, 16, 466–471.

3. Hata, T., Sano, Y., Sugawara, R., Matsumae, A., Kanamori, K., Shima, T. and Hoshi, T. (1956) Mitomycin, a new antibiotic from Streptomyces. I. *The Journal of Antibiotics*, Series A, 9, 141–146.

4. Tomasz, M. (1995) Mitomycin C: small, fast and deadly (but very selective). Chemistry & biology, 2, 575–579.

5. Iyer, V. and Szybalski, W. (1963) A molecular mechanism of mitomycin action: linking of complementary DNA strands. Proceedings of the National Academy of Sciences, 50, 355–362.

6. Tomasz, M., Chowdary, D., Lipman, R., Shimotakahara, S., Veiro, D., Walker, V. and Verdine, G.L. (1986) Reaction of DNA with chemically or enzymatically activated mitomycin C: isolation and structure of the major covalent adduct. Proceedings of the National Academy of Sciences, 83, 6702–6706.

7. Dronkert, M.L. and Kanaar, R. (2001) Repair of DNA interstrand cross-links. Mutat Res, 486, 217–247.

8. Wozniak, K.J. and Simmons, L.A. (2022) Bacterial DNA excision repair pathways. Nature Reviews Microbiology, 20, 465–477.

9. Lenhart, J.S., Schroeder, J.W., Walsh, B.W. and Simmons, L.A. (2012) DNA repair and genome maintenance in Bacillus subtilis. Microbiology and molecular biology reviews, 76, 530–564.

10. Warren, A.J., Maccubbin, A.E. and Hamilton, J.W. (1998) Detection of mitomycin C- DNA adducts in vivo by 32P-postlabeling: time course for formation and removal of adducts and biochemical modulation. Cancer research, 58, 453–461.

11. Wozniak, K.J. and Simmons, L.A. (2022) Bacterial DNA excision repair pathways. Nat Rev Microbiol, 20, 465–477.

12. Bharati, B.K., Gowder, M., Zheng, F., Alzoubi, K., Svetlov, V., Kamarthapu, V., Weaver, J.W., Epshtein, V., Vasilyev, N., Shen, L. et al. (2022) Crucial role and mechanism of transcription-coupled DNA repair in bacteria. Nature, 604, 152–159.

13. Smith, B.T., Grossman, A.D. and Walker, G.C. (2002) Localization of UvrA and effect of DNA damage on the chromosome of *Bacillus subtilis*. J. Bacteriol., 184, 488–493.

14. Kraithong, T., Hartley, S., Jeruzalmi, D. and Pakotiprapha, D. (2021) A Peek Inside the Machines of Bacterial Nucleotide Excision Repair. Int J Mol Sci, 22.

15. Dianov, G. and Lindahl, T. (1994) Reconstitution of the DNA base excision-repair pathway. Curr Biol, 4, 1069–1076.

16. Burby, P.E., Simmons, Z.W., Schroeder, J.W. and Simmons, L.A. (2018) Discovery of a dual protease mechanism that promotes DNA damage checkpoint recovery. PLoS Genetics, 14, e1007512.

17. Burby, P.E. and Simmons, L.A. (2019) A bacterial DNA repair pathway specific to a natural antibiotic. Molecular microbiology, 111, 338–353.

18. Roske, J.J., Liu, S., Loll, B., Neu, U. and Wahl, M.C. (2021) A skipping rope translocation mechanism in a widespread family of DNA repair helicases. Nucleic acids research, 49, 504–518.

19. Yang, W. (2011) Nucleases: diversity of structure, function and mechanism. Quarterly reviews of biophysics, 44, 1–93.

20. D’Andrea, L.D. and Regan, L. (2003) TPR proteins: the versatile helix. Trends in biochemical sciences, 28, 655–662.

21. Karpenahalli, M.R., Lupas, A.N. and Söding, J. (2007) TPRpred: a tool for prediction of TPR-, PPR-and SEL1-like repeats from protein sequences. BMC bioinformatics, 8, 1-8.

22. Jumper, J., Evans, R., Pritzel, A., Green, T., Figurnov, M., Ronneberger, O., Tunyasuvunakool, K., Bates, R., Žídek, A. and Potapenko, A. (2021) Highly accurate protein structure prediction with AlphaFold. Nature, 596, 583–589.

23. Liu, H. and Naismith, J.H. (2008) An efficient one-step site-directed deletion, insertion, single and multiple-site plasmid mutagenesis protocol. BMC biotechnology, 8, 1–10.

24. Gibson, D.G. (2011), Methods in enzymology. Elsevier, Vol. 498, pp. 349–361.

25. Potterton, L., Agirre, J., Ballard, C., Cowtan, K., Dodson, E., Evans, P.R., Jenkins, H.T., Keegan, R., Krissinel, E. and Stevenson, K. (2018) CCP4i2: the new graphical user interface to the CCP4 program suite. Acta Crystallographica Section D: Structural Biology, 74, 68–84.

26. McCoy, A.J., Grosse-Kunstleve, R.W., Adams, P.D., Winn, M.D., Storoni, L.C. and Read, R.J. (2007) Phaser crystallographic software. Journal of applied crystallography, 40, 658–674.

27. Emsley, P. and Cowtan, K. (2004) Coot: model-building tools for molecular graphics. Acta crystallographica section D: biological crystallography, 60, 2126–2132.

28. Murshudov, G.N., Skubák, P., Lebedev, A.A., Pannu, N.S., Steiner, R.A., Nicholls, R.A., Winn, M.D., Long, F. and Vagin, A.A. (2011) REFMAC5 for the refinement of macromolecular crystal structures. Acta Crystallographica Section D: Biological Crystallography, 67, 355–367.

29. Kenji Ohgane, H.Y. (2019.). protocols.io 10.17504/protocols.io.7vghn3w.

30. Wyrzykowska, P., Rogers, S. and Chahwan, R. (2021) Measuring Real-time DNA/RNA Nuclease Activity through Fluorescence. Bio-protocol, 11, e4206–e4206.

31. Kliegman, J.I., Griner, S.L., Helmann, J.D., Brennan, R.G. and Glasfeld, A. (2006) Structural basis for the metal-selective activation of the manganese transport regulator of Bacillus subtilis. Biochemistry, 45, 3493–3505.

32. Froschauer, E.M., Kolisek, M., Dieterich, F., Schweigel, M. and Schweyen, R.J. (2004) Fluorescence measurements of free [Mg2+] by use of mag-fura 2 in Salmonella enterica. FEMS Microbiol Lett, 237, 49–55.

33. Wakeman, C.A., Goodson, J.R., Zacharia, V.M. and Winkler, W.C. (2014) Assessment of the requirements for magnesium transporters in Bacillus subtilis. J Bacteriol, 196, 1206–1214.

34. Brucet, M., Querol-Audí, J., Serra, M., Ramirez-Espain, X., Bertlik, K., Ruiz, L., Lloberas, J., Macias, M.J., Fita, I. and Celada, A. (2007) Structure of the dimeric exonuclease TREX1 in complex with DNA displays a proline-rich binding site for WW Domains. Journal of Biological Chemistry, 282, 14547–14557.

35. Beese, L.S. and Steitz, T.A. (1991) Structural basis for the 3’-5’ exonuclease activity of Escherichia coli DNA polymerase I: a two metal ion mechanism. The EMBO journal, 10, 25–33.

36. Hamdan, S., Carr, P.D., Brown, S.E., Ollis, D.L. and Dixon, N.E. (2002) Structural basis for proofreading during replication of the Escherichia coli chromosome. Structure, 10, 535–546.

37. Yang, J. and Zhang, Y. (2015) I-TASSER server: new development for protein structure and function predictions. Nucleic acids research, 43, W174–W181.

38. 38. Waterhouse, A., Bertoni, M., Bienert, S., Studer, G., Tauriello, G., Gumienny, R., Heer, F.T., de Beer, T.A.P., Rempfer, C. and Bordoli, L. (2018) SWISS-MODEL: homology modelling of protein structures and complexes. Nucleic acids research, 46, W296–W303.

39. Beese, L.S., Derbyshire, V. and Steitz, T.A. (1993) Structure of DNA polymerase I Klenow fragment bound to duplex DNA. Science, 260, 352–355.

40. Wang, T., Sun, H.-L., Cheng, F., Zhang, X.-E., Bi, L. and Jiang, T. (2013) Recognition and processing of double-stranded DNA by ExoX, a distributive 3’–5’ exonuclease. Nucleic acids research, 41, 7556–7565.

41. Cheng, H.-L., Lin, C.-T., Huang, K.-W., Wang, S., Lin, Y.-T., Toh, S.-I. and Hsiao, Y.-Y. (2018) Structural insights into the duplex DNA processing of TREX2. Nucleic acids research, 46, 12166–12176.

42. Huang, K.-W., Liu, T.-C., Liang, R.-Y., Chu, L.-Y., Cheng, H.-L., Chu, J.-W. and Hsiao, Y.-Y. (2018) Structural basis for overhang excision and terminal unwinding of DNA duplexes by TREX1. PLoS biology, 16, e2005653.

43. Tomasz, M., Chowdary, D., Lipman, R., Shimotakahara, S., Veiro, D., Walker, V. and Verdine, G.L. (1986) Reaction of DNA with chemically or enzymatically activated mitomycin C: isolation and structure of the major covalent adduct. Proc Natl Acad Sci U S A, 83, 6702–6706.

44. Bernad, A., Blanco, L., Lázaro, J., Martin, G. and Salas, M. (1989) A conserved 3’→ 5’ exonuclease active site in prokaryotic and eukaryotic DNA polymerases. Cell, 59, 219–228.

45. Lam, W.-C., Thompson, E.H., Potapova, O., Sun, X.C., Joyce, C.M. and Millar, D.P. (2002) 3’-5’ Exonuclease of Klenow Fragment: Role of Amino Acid Residues within the Single-Stranded DNA Binding Region in Exonucleolysis and Duplex DNA Melting. Biochemistry, 41, 3943–3951.

46. Zhou, W., Richmond-Buccola, D., Wang, Q. and Kranzusch, P.J. (2022) Structural basis of human TREX1 DNA degradation and autoimmune disease. Nature communications, 13, 4277.

47. Breyer, W.A. and Matthews, B.W. (2000) Structure of Escherichia coli exonuclease I suggests how processivity is achieved. Nature structural biology, 7, 1125–1128.

48. Beese, L.S., Friedman, J.M. and Steitz, T.A. (1993) Crystal structures of the Klenow fragment of DNA polymerase I complexed with deoxynucleoside triphosphate and pyrophosphate. Biochemistry, 32, 14095–14101.

49. Hsiao, Y.-Y., Yang, C.-C., Lin, C.L., Lin, J.L., Duh, Y. and Yuan, H.S. (2011) Structural basis for RNA trimming by RNase T in stable RNA 3’-end maturation. Nature chemical biology, 7, 236–243.

50. Lormand, J.D., Kim, S.-K., Walters-Marrah, G.A., Brownfield, B.A., Fromme, J.C., Winkler, W.C., Goodson, J.R., Lee, V.T. and Sondermann, H. (2021) Structural characterization of NrnC identifies unifying features of dinucleases. Elife, 10, e70146.

51. Derbyshire, V., Grindley, N. and Joyce, C. (1991) The 3’-5’ exonuclease of DNA polymerase I of Escherichia coli: contribution of each amino acid at the active site to the reaction. The EMBO Journal, 10, 17–24.

52. Kim, S.-K., Lormand, J.D., Weiss, C.A., Eger, K.A., Turdiev, H., Turdiev, A., Winkler, W.C., Sondermann, H. and Lee, V.T. (2019) A dedicated diribonuclease resolves a key bottleneck for the terminal step of RNA degradation. Elife, 8, e46313.

53. Holm, L., Laiho, A., Törönen, P. and Salgado, M. (2023) DALI shines a light on remote homologs: One hundred discoveries. Protein Science, 32, e4519.

54. Gouge, J., Ralec, C., Henneke, G. and Delarue, M. (2012) Molecular recognition of canonical and deaminated bases by P. abyssi family B DNA polymerase. Journal of molecular biology, 423, 315–336.

55. He, Q., Lin, X., Chavez, B.L., Agrawal, S., Lusk, B.L. and Lim, C.J. (2022) Structures of the human CST-Polα–primase complex bound to telomere templates. Nature, 608, 826–832.

56. Kilkenny, M.L., Veale, C.E., Guppy, A., Hardwick, S.W., Chirgadze, D.Y., Rzechorzek, N.J., Maman, J.D. and Pellegrini, L. (2022) Structural basis for the interaction of SARS-CoV- 2 virulence factor nsp1 with DNA polymerase α–primase. Protein Science, 31, 333–344.

57. Yuan, Z., Georgescu, R., Schauer, G.D., O’Donnell, M.E. and Li, H. (2020) Structure of the polymerase ε holoenzyme and atomic model of the leading strand replisome. Nature communications, 11, 3156.

58. 58. Stokes, K., Winczura, A., Song, B., De Piccoli, G. and Grabarczyk, D.B. (2020) Ctf18-RFC and DNA Pol ɛ form a stable leading strand polymerase/clamp loader complex required for normal and perturbed DNA replication. Nucleic acids research, 48, 8128–8145.

59. Das, A.K., Cohen, P.W. and Barford, D. (1998) The structure of the tetratricopeptide repeats of protein phosphatase 5: implications for TPR-mediated protein-protein interactions. EMBO J, 17, 1192–1199.

