## Supplemental figures, tables and methods for "Structural and biochemical characterization of the mitomycin C repair exonuclease MrfB"

Kelly A. Manthei[1], Lia M. Munson[1], Jayakrishnan Nandakumar[1], Lyle A. Simmons[1*]
[1]Department of Molecular, Cellular, and Developmental Biology, University of Michigan, Ann Arbor, Michigan, USA

*Plasmid construction.*

Variant plasmids for MrfB purification were constructed with site directed mutagenesis using pPB97 (10×His-Smt3-MrfB) as a template and the appropriate primers as listed in Table S1 (1,2). Plasmid pKM228 (10×His-Smt3-MrfB-RRKR) used pKM225 (10×His-Smt3-MrfB-R212A) as a template. Truncations were constructed with Gibson assembly using two PCR products: the vector was amplified with oKM218/oKM219 and inserts with oKM240 (ΔN32-forward) and the following reverse primers oKM223 (to aa413, creating pKM218), oKM225 (to aa316, creating pKM219), and oKM239 (to aa279, creating pKM220) (3).

The plasmids for spot titers were constructed via Gibson assembly as detailed below (3,4) except for pKM233-R212A which was cloned with site directed mutagenesis using plasmid pPB110 and primers oKM249 and oKM250 (2).

pKM230-L113A: The plasmid was constructed with five PCR products: 1) the vector pPB110 was amplified using oPEB116/117; 2) the upstream portion of *amyE* and the *Pxyl* promoter were amplified using oPEB370/383; 3) the chloramphenicol resistance cassette and the downstream portion of *amyE* was amplified using oPEB557/377; 4) the 5 portion of the *mrfB-L113A* ORF was amplified using oPEB564/oKM234; and 5) the 3 portion of the *mrfB-L113A* ORF was amplified using oKM226/oPEB565. Clones were verified via Sanger sequencing using oPEB345 and oPEB348.

pKM231-R203A: The plasmid was constructed with five PCR products using the same first three products above for pKM230 in addition to: 4) the 5 portion of the *mrfB-R203A* ORF was amplified using oPEB564/oKM244; and 5) the 3 portion of the *mrfB-R203A* ORF was amplified using oKM243/oPEB565. Clones were verified via Sanger sequencing using oPEB345 and oPEB348.

pKM233-RRKR: The plasmid was constructed with five PCR products using the same first three products above for pKM230 in addition to: 4) the 5 portion of the *mrfB-RRKR* ORF was amplified using oPEB564/oKM276 with template pKM232; and 5) the 3 portion of the *mrfB-RRKR* ORF was amplified using oKM274/oPEB565 with template pKM232. Clones were verified via Sanger sequencing using oPEB345 and oPEB348.

*Strain construction.*

Strains were constructed with double crossover recombination (4). The construction of PEB318 was previously published (1).

KM4 and KM12 (*ΔmrfB, amyE::Pxyl-mrfB*): PEB318 was transformed with pPB110 digested with the restriction enzymes ScaI and KpnI. Replacement of *amyE* with *Pxyl-mrfB* by double crossover recombination was verified by testing for an inability to utilize starch and by colony PCR with primers oPEB345 and oPEB348.

KM7 and KM13(*ΔmrfB, amyE::Pxyl-mrfB-L113A*): PEB318 was transformed with pKM230 digested with the restriction enzymes ScaI and KpnI. Replacement of *amyE* with *Pxyl-mrfB-L113A* by double crossover recombination was verified by testing for an inability to utilize starch and by colony PCR with primers oPEB345 and oPEB348.

KM15 and KM16 (*ΔmrfB, amyE::Pxyl-mrfB-R203A*): PEB318 was transformed with pKM231 digested with the restriction enzymes ScaI and KpnI. Replacement of *amyE* with *Pxyl-mrfB-R203A* by double crossover recombination was verified by testing for an inability to utilize starch and by colony PCR with primers oPEB345 and oPEB348.

KM8 and KM17 (*ΔmrfB, amyE::Pxyl-mrfB-R212A*): PEB318 was transformed with pKM232 digested with the restriction enzymes ScaI and KpnI. Replacement of *amyE* with *Pxyl-mrfB-R212A* by double crossover recombination was verified by testing for an inability to utilize starch and by colony PCR with primers oPEB345 and oPEB348.

KM10 and KM18 (*ΔmrfB, amyE::Pxyl-mrfB-RRKR*): PEB318 was transformed with pKM233 digested with the restriction enzymes ScaI and KpnI. Replacement of *amyE* with *Pxyl-mrfB-RRKR* by double crossover recombination was verified by testing for an inability to utilize starch and by colony PCR with primers oPEB345 and oPEB348.

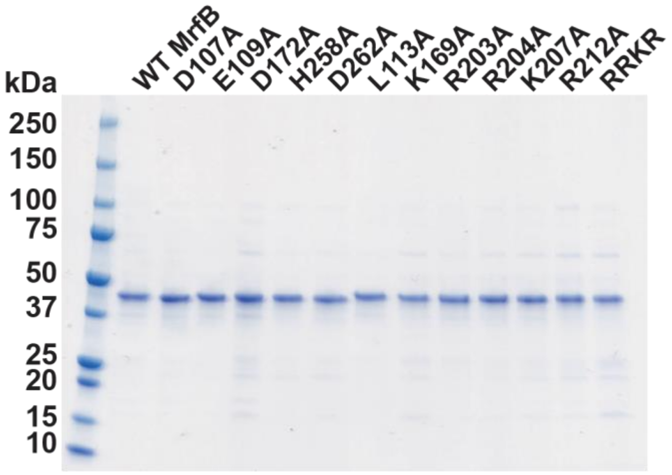

**Figure S1. MrfB variant purified protein preparations.** Approximately 1 µg was loaded to a 4-20% SDS-PAGE gel for each variant.

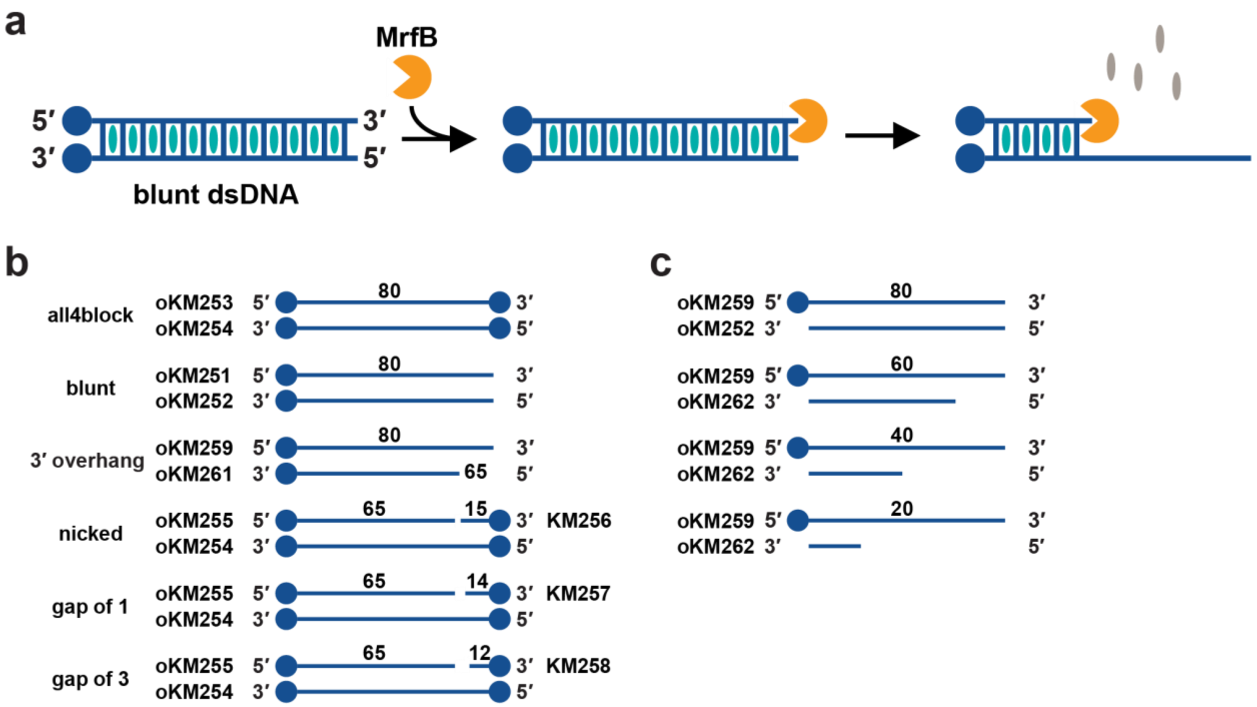

**Figure S2. PicoGreen assay schematic and substrates.** (a) Assay schematic showing that PicoGreen fluorescence starts high and decreases as MrfB acts on the DNA. The circles represent streptavidin to block the ends of DNA. (b) Various substrates used and the oligos annealed to make them. (c) DNA used for standard curve.

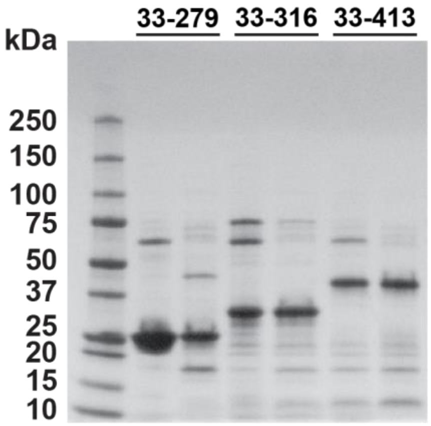

**Figure S3. SDS-PAGE gel for test expressions of MrfB truncations.** The two lanes for each construct were from the second Ni-NTA column after the His-SUMO was removed, showing the flow through (lane 1) and 40 mM imidazole wash (lane 2). The same volume was loaded for each prep, which reveals higher expression levels for the exonuclease core (residues 33-279, lane 1).

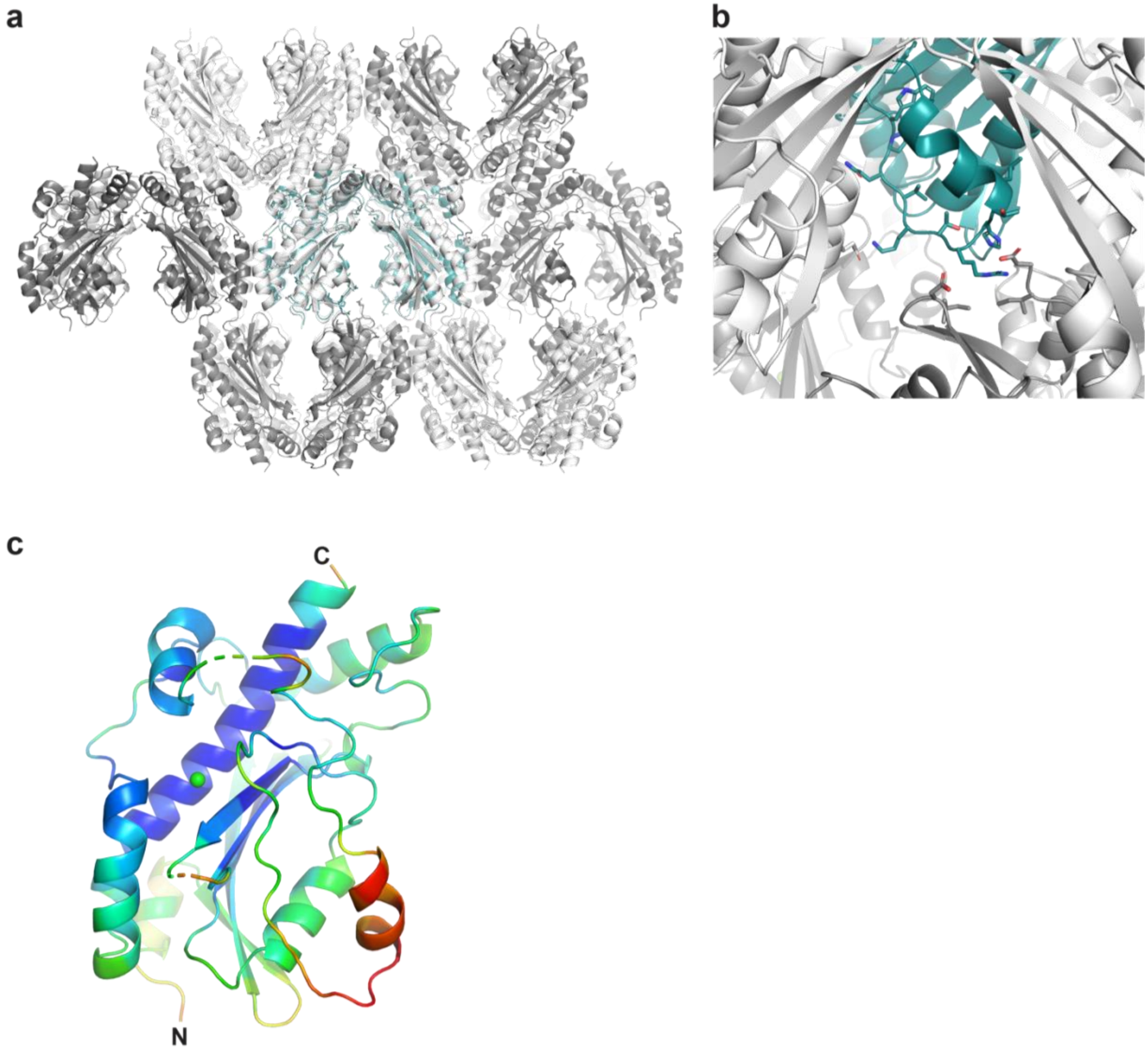

**Figure S4. Crystal packing and B-factors of the MrfB exonuclease core.** (a) Multiple symmetry mates are shown to visualize the crystal packing of the structure, with the MrfB exonuclease core asymmetric unit in teal. (b) The B chain of MrfB near the expected α-helix from 171-183 to show how it packs together. There are potential interactions but the density is not very strong for sidechains in this area and there is room for solvent in much of the region. (c) B-factors modeled onto the exonuclease cores structure. Lower B-factors are blue to green, higher B-factors are from yellow to red (highest).

**Table S1. Primers and DNA Substrate Sequences**

| Cloning Primers | Sequence |
|---|---|
| oPEB345-SeqF | ACTCCTTTGTTTATCCACCGAAC |
| oPEB348-SeqR | TTATTTTTGACACCAGACCAACTG |
| oPEB731-D107A-F | ACAAAAACAACCTCTTTTTCTTTGCTACAGAAACAACCGGTCTTGGGGGT |
| oPEB732-D107A-R | CCCCAAGACCGGTTGTTTCTGTAGCAAAGAAAAAGAGGTTGTTTTTGTTA TACCCT |
| oPEB733-E109A-F | CAACCTCTTTTTCTTTGATACAGCTACAACCGGTCTTGGGGGTGGA |
| oPEB734-E109A-R | CTCCACCCCCAAGACCGGTTGTAGCTGTATCAAAGAAAAAGAGGTTGTT TTTGTT |
| oPEB765-D262A-F | TGTCCTGCATCATAATGAAATGGCTGTGTTATCACTCATTTCATTGTACAT C |
| oPEB766-D262A-R | ACAATGAAATGAGTGATAACACAGCCATTTCATTATGATGCAGGACACCT |
| oPEB767-H258A-F | TCTTTTAAAAGGTGTCCTGCATGCTAATGAAATGGATGTGTTATCACTCA TTTC |
| oPEB768-H258A-R | GTGATAACACATCCATTTCATTAGCATGCAGGACACCTTTTAAAAGATCC |
| oKM213-D172A-F | AAAGCCTTTGCTTGGCCGCAG |
| oKM214-D172A-R | GCCGTTGTAGGTCACAAG |
| oKM235-L113A-F | AACCGGTGCTGGGGGTGGAGCTGGGAATACCATTTTTTTG |
| oKM236-L113A-R | CACCCCCAGCACCGGTTGTTTCTGTATCAAAGAAAAAGAGG |
| oKM241-F171A-F | CAAAGCCGCTGATTGGCCGCAGGTGAAAACAAGGCAC |
| oKM242-F171A-R | GCCAATCAGCGGCTTTGCCGTTGTAGGTCACAAGTGATG |
| oKM243 R203A-F | TGGAGCTGCACGCCTGTGGAAACACAAAATGGACCGTG |
| oKM244-R203A-R | CAGGCGTGCAGCTCCATGCAAAAGATCAAAGTGGCCAAAC |
| oKM245-R204A-F | AGCTAGAGCCCTGTGGAAACACAAAATGGACCGTGTATC |
| oKM246-R204A-R | CCACAGGGCTCTAGCTCCATGCAAAAGATCAAAGTGGC |
| oKM247-K207A-F | CCTGTGGGCACACAAAATGGACCGTGTATCTCTTGGC |
| oKM248-K207A-R | CTTTTGCATGGAGCTAGACGCCTGTGGGCACACAAAA |
| oKM249-R212A-F | AATGGACGCTGTATCTCTTGGCACGGTTGAAAAAGAG |
| oKM250-R212A-R | GAGATACAGCGTCCATTTTGTGTTTCCACAGGCGTC |
| oKM272-K169A-F | CAACGGCGCAGCCTTTGATTGGCCGCAGGTGAAAAC |
| oKM273-K169A-R | AAAGGCTGCGCCGTTGTAGGTCACAAGTGATGTAATGTC |
| oKM274-RRKR-F | GAGCTGCAGCCCTGTGGGCACACAAAATGGACGCTGTATC |
| oKM275-RRKR-R | CACAGGGCTGCAGCTCCATGCAAAAGATCAAAGTGG |
| oKM276-RRKR-R-Gib | CAGCGTCCATTTTGTGTGCCCACAGGGCTGCAGCTCCATGC |
| oKM240-ΔN32-F | TCACCGCGAACAGATTGGAGGTGATGACATCCCGTTTTTAGAAG |
| oKM223-aa413-R | GTGGTGGTGCTCGATTATTAAGAGGAATATTTCCTCTTTAGCCGG |
| oKM225-aa316-R | GTGGTGGTGCTCGATTATTATTTTTCTATAAGCCTTTCCAACTGCTTG |
| oKM239-aa279-R | TGGTGGTGGTGCTCGATTATTATGAAAGGATTTTTTTAGACATATG |

| **Urea-PAGE Substrates** | |
|---|---|
| oKM228 | /5IRD800/AGTAGTGAACCATGCTTACG |
| oKM229 | /5IRD800CWN/AGTAGTGAArCrCrArUrGrCrUrUrArCrG |
| oJR348 | AGTAGTGAACCATGCTTACG/3IR800CWN/ |
| **PicoGreen Substrates** | |
| oKM251-80nt-F | GATGGTTTGTTGGTCTATTACTACTTGGAGCTTGTATGATTCGAAACCTT GGAGTACTTGCCTACTTGGAGTGAACTTAG |
| oKM252-80nt-R | CTAAGTTCACTCCAAGTAGGCAAGTACTCCAAGGTTTCGAATCATACAAG CTCCAAGTAGTAATAGACCAACAAACCATC |
| oKM253-80nt-F-block | /5BiotinTEG/GATGGTTTGTTGGTCTATTACTACTTGGAGCTTGTATGATTC GAAACCTTGGAGTACTTGCCTACTTGGAGTGAACTTAG/3BioTEG/ |
| oKM254-80nt-R-block | /5BiotinTEG/CTAAGTTCACTCCAAGTAGGCAAGTACTCCAAGGTTTCGAA TCATACAAGCTCCAAGTAGTAATAGACCAACAAACCATC/3BioTEG/ |
| oKM255-65nt-F-5'block | /5BiotinTEG/GATGGTTTGTTGGTCTATTACTACTTGGAGCTTGTATGATTC GAAACCTTGGAGTACTTGCCTAC |
| oKM256-15nt-F-3'block | TTGGAGTGAACTTAG/3BioTEG/ |
| oKM25714nt-F-3'block | TGGAGTGAACTTAG/3BioTEG/ |
| oKM258-12nt-F-3'block | GAGTGAACTTAG/3BioTEG/ |
| oKM259-80nt-F-5'block | /5BiotinTEG/GATGGTTTGTTGGTCTATTACTACTTGGAGCTTGTATGATTC GAAACCTTGGAGTACTTGCCTACTTGGAGTGAACTTAG |
| oKM260-80nt-R-3'block | CTAAGTTCACTCCAAGTAGGCAAGTACTCCAAGGTTTCGAATCATACAAG CTCCAAGTAGTAATAGACCAACAAACCATC/3BioTEG/ |
| oKM261-65nt-R-3'block | GTAGGCAAGTACTCCAAGGTTTCGAATCATACAAGCTCCAAGTAGTAATA GACCAACAAACCATC/3BioTEG/ |
| oKM262-60nt-R | CAAGTACTCCAAGGTTTCGAATCATACAAGCTCCAAGTAGTAATAGACCA ACAAACCATC |
| oKM263-40nt-R | ATCATACAAGCTCCAAGTAGTAATAGACCAACAAACCATC |
| oKM264-20nt-R | TAATAGACCAACAAACCATC |

**Table S2. Strains**

| Strain | Genotype | Plasmid or gDNA | Reference |
|---|---|---|---|
| PY79 | Prototroph, Spβ° | | (5) |
| PEB318 | Δ*mrfB* | PY79 | (4) |
| PEB345 | *amyE::Pxyl-mrfB* | pPB110 | (4) |
| PEB348 | Δ*mrfB, amyE::*$P_{xyl}$*-mrfB* | pPB110 | (1) |
| PEB371 | Δ*mrfB, amyE::*$P_{xyl}$*-mrfB* | gDNA from PEB345 | (1) |
| KM4/12 | Δ*mrfB, amyE::*$P_{xyl}$*-mrfB* | pPB110 | this study |
| KM7/13 | Δ*mrfB, amyE::*$P_{xyl}$*-mrfB-L113A* | pKM230 | this study |
| KM15/16 | Δ*mrfB, amyE::*$P_{xyl}$*-mrfB-R203A* | pKM231 | this study |
| KM8/17 | Δ*mrfB, amyE::*$P_{xyl}$*-mrfB-R212A* | pKM232 | this study |
| KM10/18 | Δ*mrfB, amyE::*$P_{xyl}$*-mrfB-RRKR* | pKM233 | this study |
| BL21(DE3) | *E. coli: fhuA2[lon] ompT gal(λDE3)[dcm] ΔhsdS; λDE3= λ sBamHIo ΔEcoRI-B int::(lacI::PlacUV5::T7 gene 1) i21 Δnin5* | | Lab Stock |

### SUPPLEMENTAL REFERENCES

1. Burby, P.E. and Simmons, L.A. (2019) A bacterial DNA repair pathway specific to a natural antibiotic. *Molecular microbiology*, **111**, 338-353.
2. Liu, H. and Naismith, J.H. (2008) An efficient one-step site-directed deletion, insertion, single and multiple-site plasmid mutagenesis protocol. *BMC biotechnology*, **8**, 1-10.
3. Gibson, D.G. (2011), *Methods in enzymology*. Elsevier, Vol. 498, pp. 349-361.
4. Burby, P.E., Simmons, Z.W., Schroeder, J.W. and Simmons, L.A. (2018) Discovery of a dual protease mechanism that promotes DNA damage checkpoint recovery. *PLoS Genetics*, **14**, e1007512.
5. Youngman, P., Perkins, J.B. and Losick, R. (1984) A novel method for the rapid cloning in Escherichia coli of Bacillus subtilis chromosomal DNA adjacent to Tn 917 insertions. *Molecular and General Genetics MGG*, **195**, 424-433.
